# Reduced prefrontal synaptic connectivity and disturbed oscillatory population dynamics in the CNTNAP2 model of autism

**DOI:** 10.1101/322388

**Authors:** Maria T. Lazaro, Jiannis Taxidis, Tristan Shuman, Iris Bachmutsky, Taruna Ikrar, Rommel Santos, Giuseppe M. Marcello, Apoorva Mylavarapu, Swasty Chandra, Allison Foreman, Rachna Goli, Duy Tran, Nikhil Sharma, Michelle Azhdam, Hongmei Dong, Olga Peñagarikano, Sotiris Masmanidis, Bence Rácz, Xiangmin Xu, Daniel H. Geschwind, Peyman Golshani

## Abstract

Loss of function mutations in CNTNAP2 cause a syndromic form of autism spectrum disorder (ASD) in humans and produce social deficits, repetitive behaviors, and seizures in mice. Yet, the functional effects of these mutations at the cellular and circuit level remain elusive. Using laser scanning photostimulation, whole-cell recordings, and electron microscopy, we found a dramatic decrease in functional excitatory and inhibitory synaptic inputs in L2/3 medial prefrontal cortex (mPFC) of Cntnap2 knock-out (KO) mice. In accordance with decreased synaptic input, KO mice displayed reduced spine and synapse densities, despite normal intrinsic excitability and dendritic complexity. To determine how this decrease in synaptic inputs alters coordination of neuronal firing patterns *in vivo*, we recorded mPFC local field potentials (LFP) and unit spiking in head-fixed mice during locomotion and rest. In KO mice, LFP power was not significantly altered at all tested frequencies, but inhibitory neurons showed delayed phase-firing and reduced phase-locking to delta and theta oscillations during locomotion. Excitatory neurons showed similar changes but only to delta oscillations. These findings suggest that profound ASD-related alterations in synaptic inputs can yield perturbed temporal coordination of cortical ensembles.

## INTRODUCTION

Autism Spectrum Disorder (ASD) is characterized by deficits in social communication and repetitive or restrictive behaviors^1^. Genetic studies have revealed that the etiology of ASD is very heterogeneous, involving many hundreds of genes^2–5^, a significant proportion of which appear as rare recessive or de novo dominant mutations^6–9^. One highly penetrant syndromic form of ASD is caused by loss-of-function mutations in the CNTNAP2 gene^10^ and CNTNAP2 polymorphisms have been associated with increased risk for ASD and other conditions such as language impairments, schizophrenia, hyperactivity, and alterations in functional brain connectivity^11–13^.

CNTNAP2 encodes for Contactin-associated protein-like 2 (Caspr2), a protein of the neurexin superfamily that has diverse cellular and circuit functions, and mediates neuron-glia interactions^14^, clustering of potassium channels^15,16^, stabilization of dendritic spines^17^, glutamatergic receptor localization^18^, and development of inhibitory circuits^10,19–21^. Mice lacking the Cntnap2 gene recapitulate the core behavioral deficits of ASD, including impairments in socialization, communication, and repetitive behaviors, and display seizures^20^. Recent *in vivo* evidence suggests that CNTNAP2 has a putative role in synapse formation and stabilization and that dendritic spine dynamics are affected in the Cntnap2 KO mice, with reduced stability in newly formed spines^17^. In addition, the loss of CNTNAP2 leads to synaptic alterations *in vitro* with decreased inhibition and axonal excitability deficits in acute hippocampal slices^21–23^. These results suggest that CNTNAP2 mutations may be linked to abnormal behavior by altering synaptic neurotransmission, functional connectivity, and neuronal network activity. However, the specific cellular and circuit mechanisms that lead to altered behavior in Cntnap2 KO mice remain unclear.

Here, we examined the neurophysiological consequences of Cntnap2 deletion in the mouse medial prefrontal cortex (mPFC), a brain region critically involved in social behavior^24,25^ and notably affected in ASD^26–28^. Using glutamate uncaging via laser-scanning photostimulation (LSPS) on L2/3 pyramidal neurons of the mPFC, we observed a reduction in both excitatory and inhibitory synaptic inputs onto excitatory neurons. Through *in vitro* whole-cell patch clamp recordings, we confirmed that absence of Cntnap2 leads to decreased excitatory neurotransmission, with a concomitant decrease in dendritic spine and synapse densities. Finally, we recorded single units from excitatory and inhibitory neurons and local field potentials (LFP) in mPFC *in vivo* using multichannel silicon microprobes. We observed robust alterations in the phase-locking of units to delta and theta oscillations during locomotion, but not during rest.

These findings demonstrate that CNTNAP2 mutations result in decreased excitatory drive onto single cells, which leads to alterations in circuit-level synchronous activity in mPFC, a brain area involved in modulating social behavior. Linking CNTNAP2 mutations with specific microcircuit alterations and abnormal neural activity provides new insights into the relationship between genetic, circuit, and behavioral abnormalities in ASD.

## RESULTS

**Laser scanning photostimulation (LSPS) reveals decreased excitatory and inhibitory inputs in the mPFC of Cntnap2 KO mice.** To test how loss of CNTNAP2 alters mPFC microcircuits, we used LSPS via glutamate uncaging to measure local excitatory and inhibitory cortical inputs onto L2/3 mPFC pyramidal neurons. These cells can modulate social behavior, are critical for corticocortical communication, and have been implicated by transcriptomic studies as a critical hub for autism-related gene expression^25,28–30^. We mapped and quantified excitatory and inhibitory synaptic inputs by voltage-clamping patched pyramidal neurons at the membrane potentials of −70 mV and excitatory +5 mV, respectively, while uncaging glutamate and activating small clusters of surrounding neurons (Fig. 1a,b, Supplementary Fig. 1). We observed that, similar to WT, L2/3 pyramidal neurons in KO mice receive most of their excitatory and inhibitory synaptic inputs from L2/3 and L5 in mPFC (Fig. 1c,d). However, compared to WT, L2/3 excitatory neurons in KO mice display a dramatic reduction in both excitatory and inhibitory local synaptic inputs (Fig. 1c-e), while the balance of excitation to inhibition (E/I) in individual neurons is not significantly altered (Fig. 1f). Importantly, this decrease in synaptic strength is not due to lower neuronal responsiveness to glutamate uncaging in KO mice, since both mouse groups showed equivalent responses to glutamate uncaging onto perisomatic regions (Supplementary Fig. 1). These results demonstrate a robust reduction of local excitatory and inhibitory input onto layer 2/3 pyramidal cells in the mPFC of KO mice.

**Intrinsic excitability of pyramidal neurons and parvalbumin-postive inhibitory neurons**. We then examined whether loss of Cntnap2 resulted in altered excitability and intrinsic properties in cortical neurons of the mPFC, as Caspr2 has a known role in the clustering of potassium channels in the juxtaparanodes of axons, important for the propagation of action potentials^14–16^. This, together with the fact that Cntnap2 KO mice display epileptic seizures after three months of age, and decreased neuronal synchrony *in vivo*, suggests that alterations in neuronal excitability might be at play in producing these phenotypes. Moreover, our input mapping findings could also be associated with alterations in the intrinsic excitability of cortical excitatory and inhibitory neurons. To this end, we performed whole-cell current-clamp recordings on mPFC L2/3 pyramidal and parvalbumin-positive (PV+) inhibitory neurons. We performed whole-cell current-clamp recordings on mPFC L2/3 pyramidal and PV+ inhibitory neurons (recorded in CNTNAP2-PV-Cre X Ai9 animals) in KO and WT controls. We focused on PV+ interneurons, as these cells provide powerful perisomatic inhibition to cortical pyramidal neurons and their dysfunction has been implicated autism-associated deficits resulting from loss of Cntnap2^20,23^. Input-output curves showing average number of action potentials elicited by increasing current injections in pyramidal neurons and parvalbumin-positive inhibitory neurons revealed no statistically significant alterations in action potential firing rate (Supplementary Fig. 2) between the two groups. Action potential threshold, amplitude, half-width, afterhyperpolarization (AHP) potential, or time from peak to AHP were also not significantly different between WT and KO. Moreover, resting membrane potential, input resistance, cell membrane capacitance and membrane time constant were also not significantly different (Supplementary Table 1). This indicates that loss of Cntnap2 does not affect the intrinsic excitability of L2/3 neurons of mPFC.

**Fig. 1.**
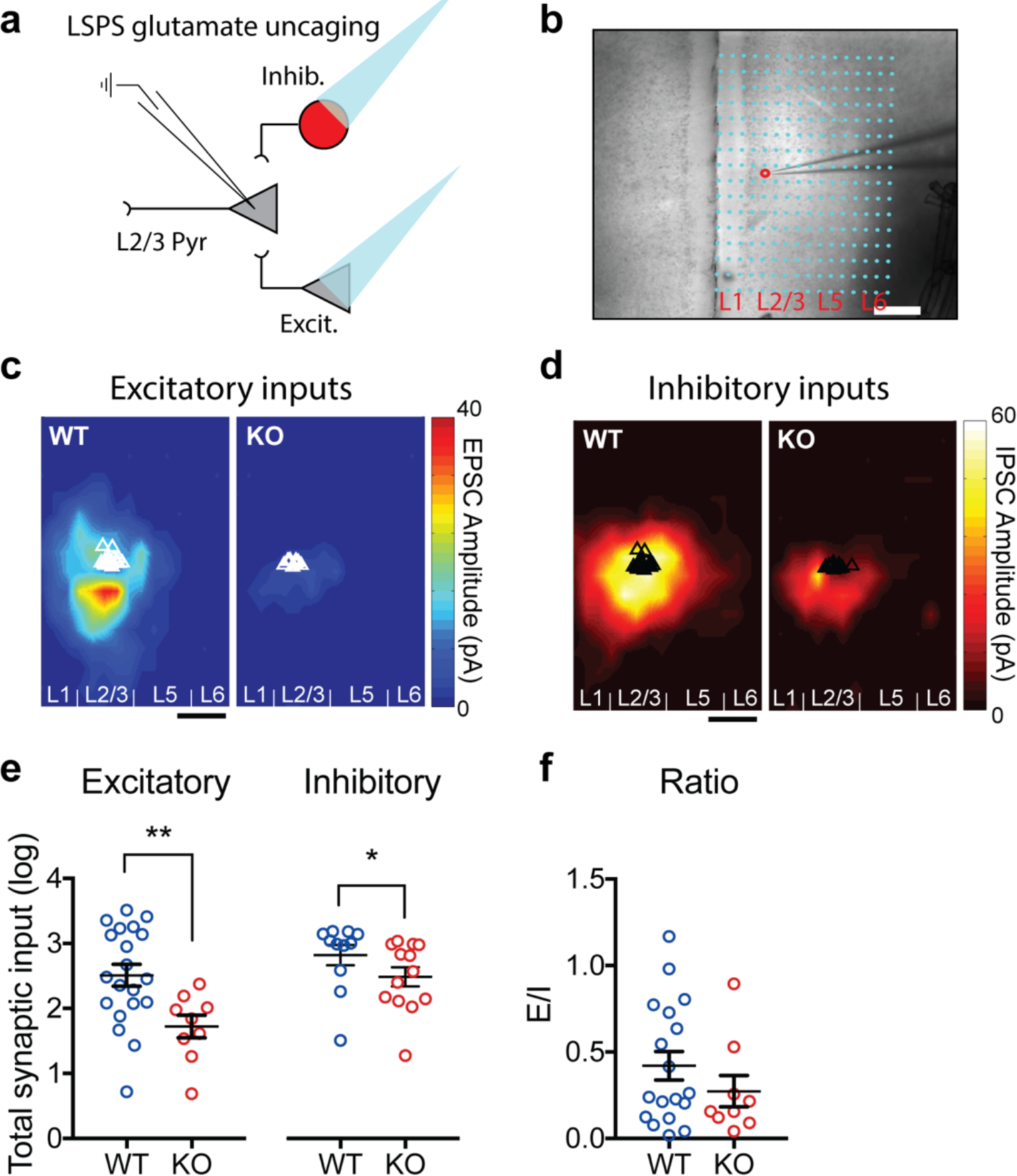
Reduced excitatory and inhibitory synaptic inputs to L2/3 pyramidal neurons in the mPFC of Cntnap2 KO mice. **a**, Schematic of experimental paradigm using laser scanning photostimulation (LSPS) via glutamate uncaging, combined with whole-cell patch clamp recordings of L2/3 pyramidal (Pyr) neurons in acute slices of the medial prefrontal cortex (mPFC). Patched neurons were clamped at −70 mV or +5 mV for detection of local excitatory or inhibitory synaptic connections, arising from photostimulated presynaptic glutamatergic (Excit.) and GABAergic (Inhib.) neurons. **b**, Example of an LSPS experiment in an mPFC slice, where differential interference contrast (DIC) imaging was used for tissue visualization. Photostimulation sites are superimposed (cyan dots) and spaced within a 100 μm × 60 μm grid. Red circle indicates the location of recorded glutamatergic neuron in L2/3, approached by the patch pipette (electrode). Scale bar represents 200 μm. **c**, Group-averaged, excitatory input maps of L2/3 excitatory neurons for WT (*n* = 20 cells, *n* = 3 mice) and KO (*n* = 9 cells, *n* = 3 mice). White triangles represent location of individually-recorded neurons. The color scale represents excitatory input strength (blue = low, red = high). Scale bar is 200 μm. **d**, Group-averaged, inhibitory input maps of L2/3 excitatory cells for WT (*n* = 11 cells, *n* = 3 mice) and KO (*n* = 13 cells, *n* = 3 mice). Black triangles represent individual recorded neurons. The color scale represents inhibitory input strength (black = low, white = high). Scale bar is 200 μm. **e**, Average total synaptic excitatory and inhibitory input strength (log) measured for L2/3 excitatory cells depicting a robust decrease in the KO, compared to WT (WT EPSC 2.51 ± 0.17, *n* = 20 cells; KO EPSC 1.72 ± 0.17, *n* = 9 cells; WT IPSC 2.83 ± 0.16, *n* = 11 cells; KO IPSC 2.49 ± 0.15, *n* = 13 cells; EPSC: \*\**P* = 0.0051, IPSC: \**P* = 0.0218; Mann-Whitney Test). **f**, Average ratios of total excitatory inputs (excitatory postsynaptic currents, EPSC) versus total inhibitory inputs (inhibitory postsynaptic currents, IPSC) from individual cells (WT *n* = 17 cells; KO, *n* = 8 cells). There is no significant difference in E/I ratio between WT and KO (*P* = 0.8873; Unpaired t test).

**Decreased excitatory neurotransmission in pyramidal neurons of Cntnap2 KO mice.** To investigate the specific cellular processes that lead to reduced synaptic responses in KO mice, we performed whole-cell patch-clamp recordings of miniature excitatory and inhibitory postsynaptic currents (mEPSCs and mIPSCs, respectively) in mPFC L2/3 neurons. We measured mEPSC and mIPSC amplitude, frequency, and kinetics, as changes in amplitude are a reliable measure of the number of receptors at synapses (quantal size), while frequency correlates with the number of contacts or probability of release^31^. In agreement with our LSPS findings, we observed a two-fold decrease in the frequency of mEPSCs (Fig. 2a,b) and a significant decrease in the average amplitude of mEPSCs in Cntnap2 KO pyramidal neurons (Fig. 2c). Together, these findings point towards a reduction in excitatory neurotransmission in mPFC pyramidal neurons of Cntnap2 KO mice, respectively. Interestingly, we observed no statistically significant alterations in frequency, amplitude, or kinetics of mIPSC (Fig. 2d-f) despite reduced inhibitory input (Fig 1d,e), which could reflect compensatory changes between synapse number and release probability or altered distribution of proximal and distal inhibitory inputs. In addition, we examined miniature postsynaptic currents in PV+ interneurons and found no significant differences in mEPSCs (Supplementary Fig. 3a-c) or mIPSCs (Supplementary Fig. 3d-f).

**Fig. 2.**
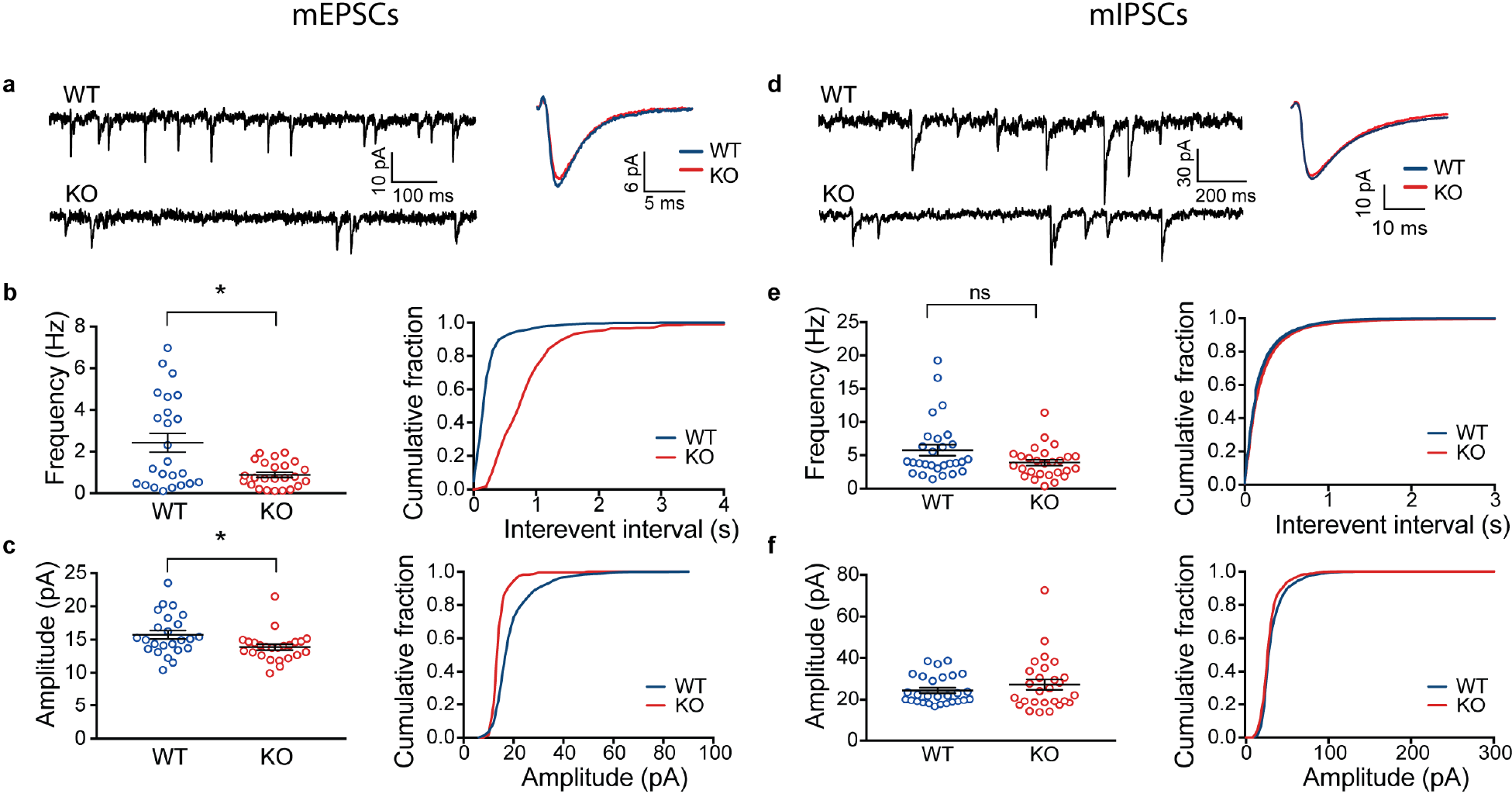
Cntnap2 KO pyramidal neurons display a two-fold decrease in the frequency of miniature excitatory postsynaptic currents (mEPSCs). **a**, Representative traces from recorded mEPSCs in Cntnap2 WT and KO pyramidal cells, voltage-clamped at −70 mV, with corresponding average unitary events. **b**, Frequency of mEPSCs (WT 2.42 ± 0.45 Hz, *n* = 24 cells; KO 0.89 ± 0.12 Hz, *n* = 24 cells; Mann-Whitney test, \**P* = 0.0410), is decreased in KO mice,. **c**, Amplitude of mEPSCs (WT 15.73 ± 0.63 pA, *n* = 24; KO 13.88 ± 0.45 pA, *n* = 24; Mann-Whitney test, \**P* = 0.0172) is decreased in KO mice. **d-f**, Same as **a-c** for miniature inhibitory postsynaptic currents (mIPSCs). There is no statistically significant decrease in frequency (WT 5.76 ± 0.83 Hz, *n* = 28; KO 3.87 ± 0.45 Hz, *n* = 27; Mann-Whitney test, *P* = 0.1327) or amplitude of mIPSCs (WT 24.43 ± 1.30 pA, *n* = 28; KO 27.21 ± 2.48 pA, *n* = 27; Mann-Whitney test, *P* = 0.8740) in KO mice compared to WT.

We then asked whether the observed decrease in mEPSC frequency on pyramidal neurons could be caused by a disruption in the probability of synaptic vesicle release^32,33,34^. We stimulated long-range axonal projections to mPFC in slices (Methods) and measured evoked excitatory currents elicited in L2/3 pyramidal cells (Fig. 3a). We observed reduced amplitude of evoked excitatory postsynaptic currents (Fig. 3b) and significantly increased EPSC latencies (Supplementary Fig. 4) in KO mice compared to controls, corroborating our previous findings of reduced excitatory neurotransmission. However, we found no significant difference in paired-pulse ratios of evoked currents between WT and KO mice (Fig. 3c), indicating similar excitatory neurotransmitter release probabilities or alterations in short-term plasticity.

We next tested whether KO mice had altered, immature or silent synapses, characterized by decreased ratio of AMPA/NMDA receptors^32,34,35,36^. We recorded evoked AMPA and NMDA currents in the presence of the GABA-A receptor blocker, picrotoxin, by holding the cells at −70 mV and at +40 mV in voltage-clamp, respectively. We found no significant difference in the AMPA-NMDA ratios when comparing WT and KO mice (Fig. 3e,f), indicating that KO mice do not have more immature or silent synapses. Together, these results indicate a reduction in the frequency and amplitude of excitatory drive onto single pyramidal cells, which cannot be explained by alterations in single synapse maturity or neurotransmitter vesicle release.

**Fig. 3.**
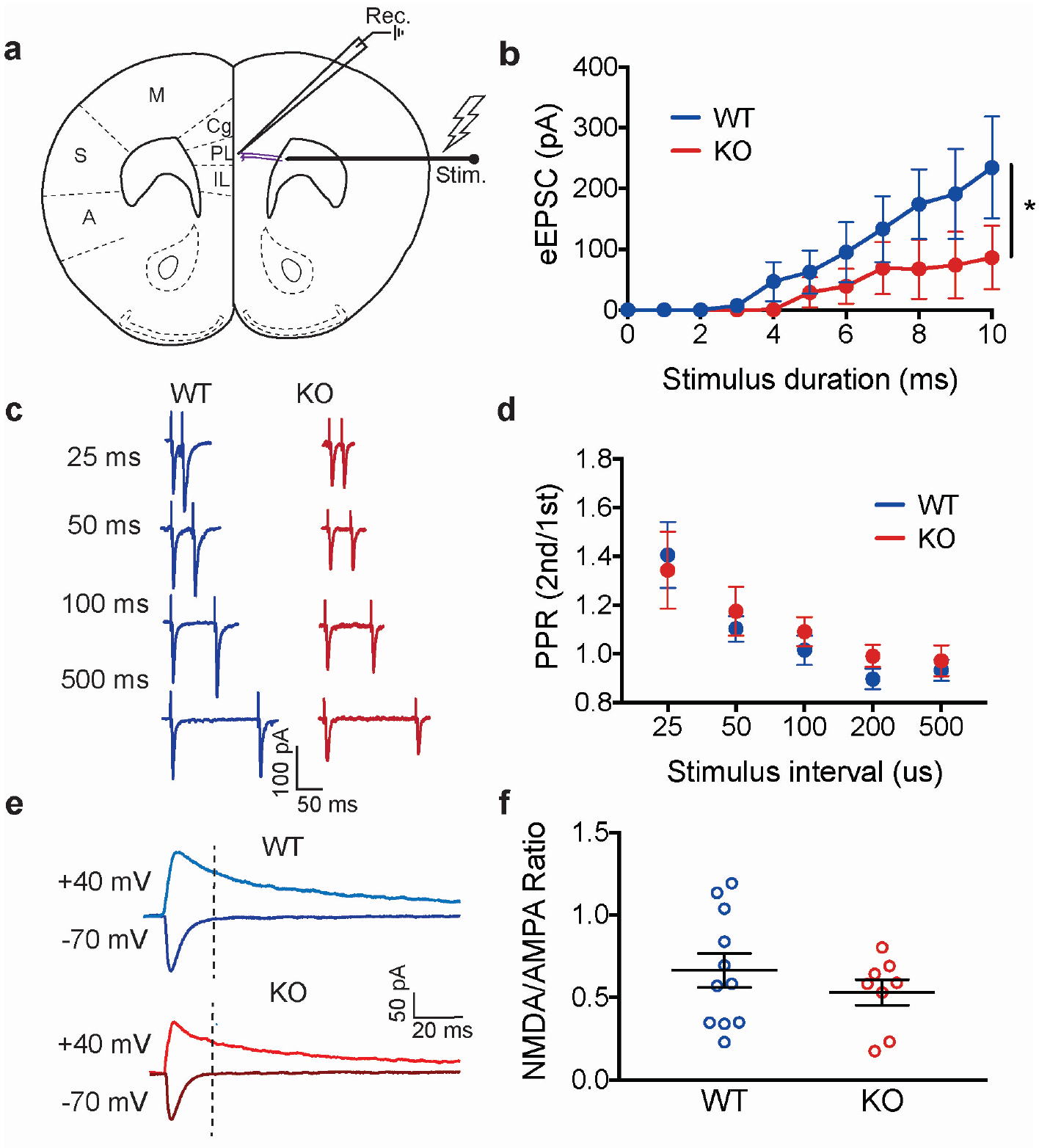
Evoked synaptic responses reveal decreased long range excitatory inputs in Cntnap2 KO mice that are not associated with altered AMPA/NMDA ratios or synaptic vesicle release. **a**, Monopolar tungsten electrode was used to stimulate long-range axons (purple), which extend from the anterior forceps of the corpus callosum and project onto a patched excitatory neuron in L2/3 mPFC. **b**, Input-output curves of unitary excitatory responses resulting from a range of increasing stimulus intensities in Cntnap2 WT and KO mice (WT *n* = 7 cells, KO *n* = 9 cells; \**P* < 0.0001, 2way ANOVA). **c**, Representative current responses from paired-pulses given at various inter-stimulus intervals (ISIs) in WT and KO mice. **d**, Ratio of 2nd /1st evoked synaptic response to paired-pulse stimulation at increasing ISIs suggests no significant deficits in the probability of synaptic vesicle release in Cntnap2 KO mice (WT *n* = 10 cells, KO *n* = 8 cells; *P* = 0.8926, 2way ANOVA). **e**, Evoked AMPA (cells voltage-clamped at −70mV) and NMDA (cells voltage-clamped at +40mV) currents in WT and KO mice. Stimulus artifact was blanked for clarity. Dashed line indicates point where NMDA current amplitudes were measured, immediately after AMPA current decay. **f**, AMPA/NMDA ratios of Cntnap2 KO mice were not significantly altered, compared to WT mice, suggesting no significant changes in synaptic maturity (WT 0.67 ± 0.10, *n* = 11 cells; KO 0.53 ± 0.08, *n* = 8 cells; *P* = 0.3471, Unpaired t test).

**Decreased dendritic spine density in Cntnap2 KO mice.** We next asked whether the decrease in excitatory neurotransmission was caused by a reduction in the total number of synaptic inputs, either through decreased dendritic branching or decreased spine density. We performed 3D anatomical reconstructions of L2/3 pyramidal neurons by filling cells with biocytin during *in vitro* slice recording experiments and imaging them with confocal microscopy. Sholl analysis did not reveal any significant changes in total dendritic length, total number of dendritic branches or in dendritic complexity (Fig. 4a,b), suggesting normal dendritic arborization in KO mice.

To determine whether Cntnap2 KO neurons display a decrease in dendritic spine density, we used Cntnap2 (homozygous KO or WT) x Thy1-GFP mice, which express GFP in a subset of pyramidal neurons, including L2/3 of mPFC (Fig. 4c). Quantification of dendritic spines of Thy1-GFP-positive pyramidal neurons in L2/3 of mPFC unveiled a significant decrease in the density of spines in both basal and apical dendritic branches (Fig. 4c,d). This suggested that the decreased functional synaptic inputs we observed are likely due to a decrease in excitatory synapse density.

To further test this hypothesis, we used electron microscopy to study L2/3 mPFC of WT (n=3) and Cntnap2 KO mice (n = 3), examining dendritic spines, and synaptic contacts (Fig. 4e). Consistently with our previous findings, we observed a significant (~25%) reduction in the number of postsynaptic spines with postsynaptic densities in KO mice (Fig. 4f). Furthermore, we found no significant changes in spine area or synapse length in KO mice (Fig. 4g). We did, however, find that KO mice had reduced density of multi-synaptic boutons (Fig. 4h), a marker of synaptogenesis which is increased following synaptic plasticity^32^. We also found an increase in perforated synapses in KO mice (Fig 4i), which are associated with increased synaptic turnover^33,34^ supporting previous reports of increased spine turnover in KO mice^17^. Together, these findings indicate that KO mice have significant defects in inhibitory and excitatory synaptic density and alterations in markers of synapse plasticity and stability.

**Fig. 4.**
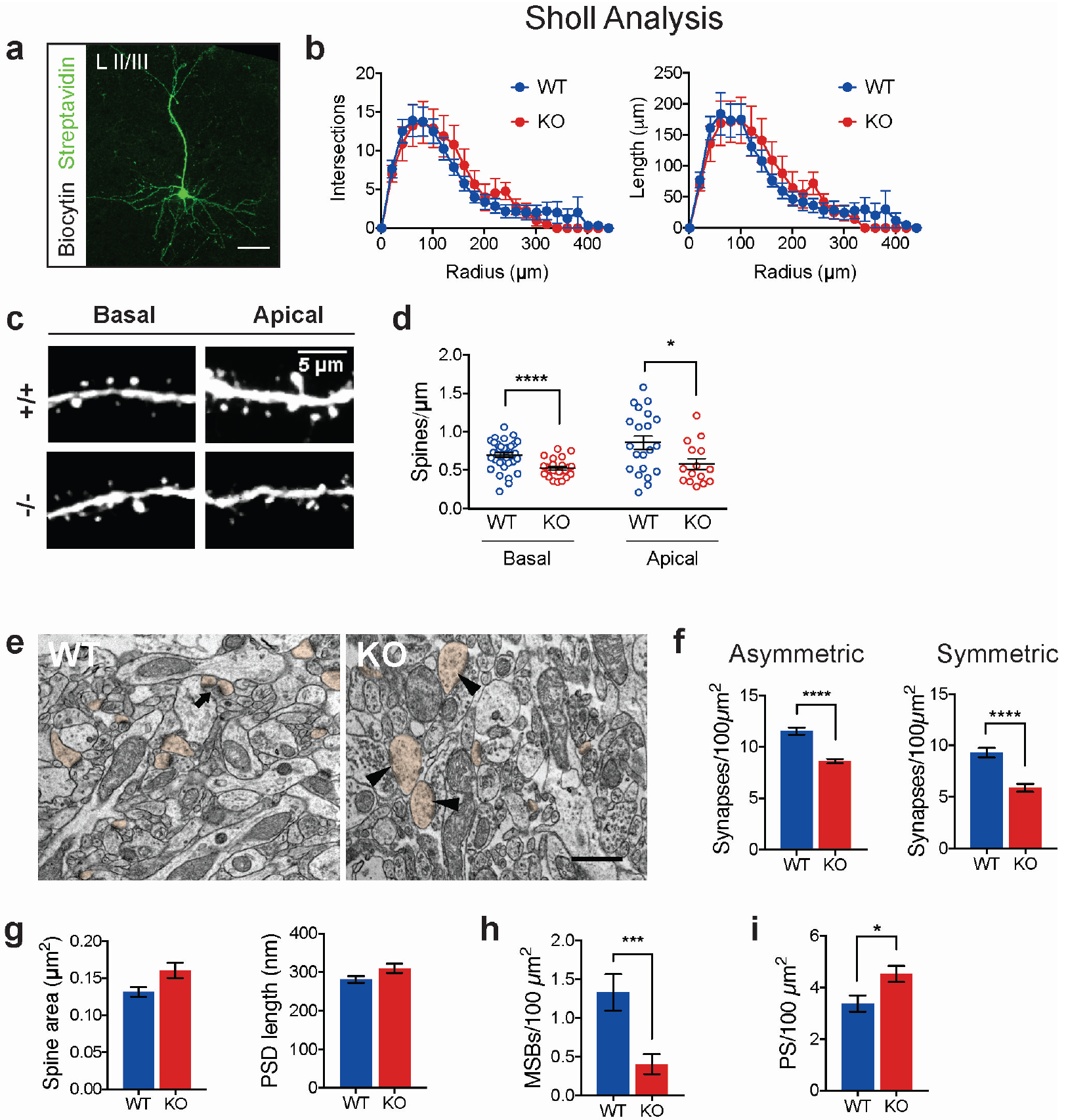
Decreased dendritic excitatory and inhibitory synapses in Cntnap2 KO mice. **a**, Representative z-stack projection of biocytin-filled L2/3 neuron, visualized with a Streptavidin ‑ Alexa 488 antibody. Scale bar indicates 100 um length. **b**, Sholl analysis showing number of intersections (*P* = 0.0632, 2-way ANOVA) and length (*P* = 0.9315, 2way ANOVA) of dendrites is comparable between Cntnap2 WT (*n* = 8 cells) and KO (*n* = 9 cells) STATS. **c**, Confocal image of L2/3 Thy1-GFP-positive pyramidal neurons in mPFC, demonstrating representative apical and basal dendrites for Cntnap2 WT (+/+) and KO (−/−) mice. **d**, Summary graphs showing quantification of average spine density in apical (WT 0.86 ± 0.09 spines/μm, *n* = 21 dendrites; KO 0.57 ± 0.07 spines/μm, *n* = 15 dendrites) and basal (WT 0.70 ± 0.03 spines/μm, *n* = 34 dendrites; KO 0.52 ± 0.02, *n* = 24 dendrites) branches. Statistical significance is represented by * for unpaired t test *P* < 0.05 and **** for Welch’s t test *P* < 0.0001. **e**, Representative electron micrographs showing neuropil of L2/3 mPFC in Cntnap2 WT and KO mice. Arrow indicates a multi-synapse bouton (MSB), arrowheads indicate perforated asymmetrical synapses. Spine profiles are pseudo-colored in orange. Scale bar: 500 nm. **f**, Graphs showing quantification of asymmetric (putative excitatory: WT 11.53 ± 0.35 synapses/100 μm^2^, *n* = 80 fields; KO 8.59 ± 0.21 synapses/100 μm^2^, *n* = 90 fields; 3 mice per genotype; \*\*\*\**P* < 0.0001, Mann-Whitney test) and symmetric (putative inhibitory: WT 9.30 ± 0.46 synapses/100 μm^2^, *n* = 80 fields; KO 5.87 ± 0.38 synapses/100 μm^2^, *n* = 90 fields; \*\*\*\**P* < 0.0001, Mann-Whitney test) synapses. **g**, Spine/postsynaptic density (PSD) area (WT 0.1313 ± 0.006 μm^2^, *n* = 257 spines; KO 0.1604 ± 0. 010 μm^2^, *n* = 187 spines; *P* = 0.1142, Mann-Whitney test) and length (WT 281.10 ± 8.89 nm, *n* = 257 spines; KO 309.90 ± 11.83 nm, *n* = 187 spines; *P* = 0.0709 Mann-Whitney test). **h**, Density of MSBs for WT (1.33 ± 0.23 MSBs per 100 μm^2^, *n* = 80 fields) and KO (0.41 ± 0.13 MSBs per 100 μm^2^, *n* = 90 fields); *** *P* = 0.0002, Mann-Whitney test. **i**, Perforated synapses (PS) for WT (3.38 ± 0.31 PS per 100 μm^2^, *n* = 90 fields) and KO (4.52 ± 0.31 PS per 100 μm^2^, *n* = 80 fields); *\*P* = 0.0122, Mann-Whitney Test.

**Altered *in vivo* network activity in mPFC of Cntnap2 KO mice.** Different cortical neuronal populations are selectively recruited to fire at specific phases of ongoing oscillations, creating a dynamic circuit pattern (or “chronocircuit”)^35^, where distinct cell types are activated in a brain-state specific manner. The precise temporal coordination of neuronal firing during oscillations is critically dependent on the balance of excitatory and inhibitory inputs. We therefore examined whether the synaptic alterations we observed in Cntnap2 KO mice had specific functional consequences on the temporal coordination of mPFC circuits during neuronal oscillations.

We recorded *in vivo* local field potentials and single unit activity in the mPFC of KO and WT mice using multi-channel silicon microprobes^36^ (Fig. 5a,b). Mice were head-fixed but free to rest or run on a spherical treadmill during the recordings^37^. Since coordinated collective synaptic activity is thought to shape the LFP signal, particularly at low frequencies^38^, we first tested if LFP low-frequency oscillations were altered in KO mice. We found no significant differences in the average LFP power between KO and WT mice in delta (1-4 Hz) or theta (5-11 Hz) oscillations during either locomotion or immobility (Supplementary Fig. 5). We found no differences in power for higher frequency band oscillations either (beta 12-30 Hz, slow gamma 30-55 Hz, or high gamma 80-110 Hz; Supplementary Fig 7), indicating no major alterations in the oscillations at the broader neuronal population level.

We also recorded 249 single units from 8 WT mice and 145 units from 5 KO mice, which were clustered into wide-spiking (WS) putative excitatory units and narrow-spiking (NS) putative interneurons (Fig. 5b, Methods). During both locomotion and immobility, firing rates of WS putative excitatory neurons were similar between the WT and KO groups (Fig. 5c, Supplementary Fig. 6). However, NS units from KO mice fired at a higher rate compared to the WT group in both states (Fig. 5c, Supplementary Fig. 6).

We next asked how the synaptic alterations we found in KO mice reflect on the spiking modulation of individual units during LFP oscillations. We examined the preferred phase of each unit’s spiking and the strength of the unit’s phase-locking to that preferred phase, focusing again on delta and theta frequency LFP oscillations (Fig. 5d,e; Supplementary Fig. 6). We found a significant decrease in the strength of phase-locking of WS units to delta oscillations and NS units to delta and theta oscillations in KO animals during locomotion (Fig. 5f,g). In addition, WS units from KO animals on average fired later in the delta oscillatory cycle, and NS units fired later in both delta and theta oscillatory cycles in KO animals (Fig. 5f,g). During immobility, changes in phase-locking to delta oscillations were not observed, whereas in theta oscillations we again found significant forward-shifted preferred phases in WS units and reduced phase-locking in NS units (Supplementary Fig 5). An overall similar trend for KO units to have forward shifted preferred phases and weaker phase-locking was observed when repeating this analysis over higher LFP oscillation frequencies as well (Supplementary Fig 6), but differences were not significant in most cases. Notably, we found a decrease in phase-locking to fast gamma oscillations in NS units of KO mice (Supplementary Fig 6), but over a small sample of phase-locked units. Finally, a similar percentage of units per animal was significantly phase-locked to each frequency in both WT and KO groups (Supplementary Fig 8).

Together, these results indicate a disrupted mPFC network in Cntnap2 KO mice, where both excitatory and inhibitory neurons have less precise firing patterns, that are also shifted relative to network activity, and point towards less coherent network dynamics. These alterations may lead to severely altered mPFC processing in Cntnap2 KO mice, potentially contributing to the altered brain function these mice.

**Fig. 5.**
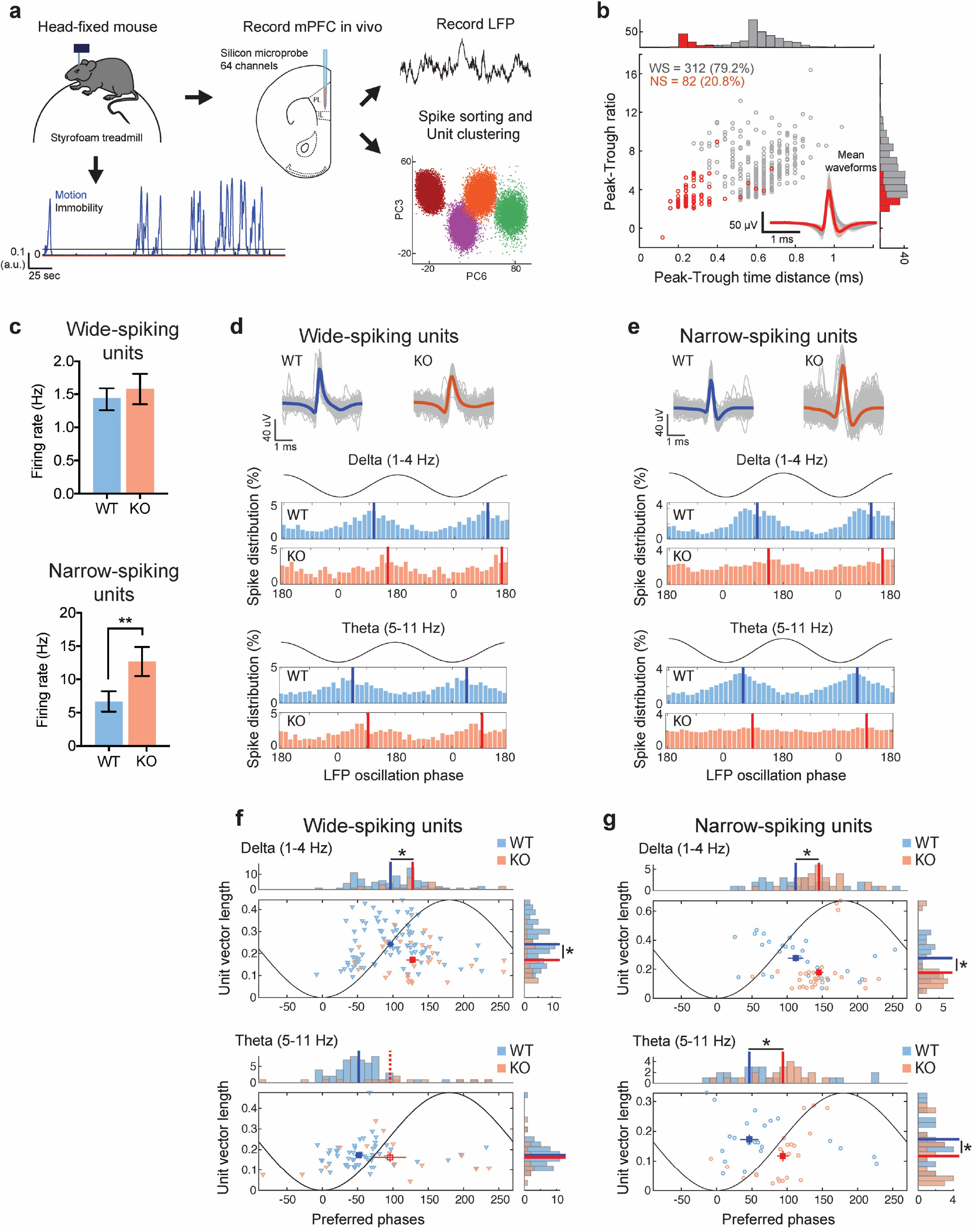
*In vivo*-recorded single units exhibit altered spiking and phase-locking to low-frequency LFP oscillations during locomotion in Cntnap2 KO mice. **a**, Schematic of *in vivo* recordings and subsequent analysis of electrophysiological data. Mice were head-fixed while free to run on a spherical treadmill. Extracellular signals were recorded from all layers of mPFC of both WT (*n* = 8) and KO mice (*n* = 5) using multi-electrode silicon microprobes. Local field potentials (LFP) and extracellular spikes were extracted and spikes were sorted into tentative single units using the PyClust software (Methods). Motion on the treadmill was also recorded and traces were separated into locomotion and immobility segments. Only concatenated locomotion segments are analyzed here. **b**, Units from all animals were split into two clusters based on their average waveform characteristics and their burst index (Methods). Distribution and corresponding histograms of mean waveform peak-to-trough ratios versus peak-to-trough time distances for the two unit-clusters. Narrow-spiking units (NS; tentative interneurons, ~20% units) are shown in red and wide-spiking units (WS; tentative pyramidal cells, ~80%) in grey. Inset depicts the average waveforms from all units in the two clusters. **c**, Average firing rate during locomotion per WS unit (top) and NS unit (bottom) in WT versus KO mice. WS units spiked with similar rates in both mouse groups, whereas NS units from KO mice fired significantly more than WT mice *P* < 0.01; Wilcoxon test). **d**, Top: Unit spike waveforms averaged over all individual spikes (gray) from two example WS units from a WT (blue) and a KO mouse (red). Middle: LFP delta frequency phase-histograms of spikes (% over all spikes) of the two units. Two cycles of the oscillation are shown together with a representation of the LFP oscillation for clarity. Solid lines indicate the mean (‘preferred’) phase of each unit. Bottom: Similar for theta frequency phase distributions. In both cases the KO unit exhibited a forward-shifted preferred phase. **e**, Same as **d** for two example NS units from a WT and a KO mouse. The KO unit’s preferred phase is again shifted and its phase-locking is weaker compared to the WT one. **f**, Distributions and corresponding distribution histograms of strength of phase-locking (mean vector length) versus preferred phases of pooled WS units from WT (blue) and KO mice (red). Only significantly phase-locked units with sufficient spiking are included (Methods). Filled rectangles and solid lines depict the mean (±SE) of the distributions and histograms respectively (circular mean for phase distributions) for the two mouse groups. All distributions were significantly non-uniform (*P* < 0.05; Rayleigh test for non-uniformity; Bonferroni corrected over frequencies and mouse groups) except for theta frequency distribution of KO units (indicated by open rectangle and dashed lines accordingly). Asterisks indicate significant differences either in mean vector length (*P* < 0.05; t-test if both distributions are normally distributed according to Lilliefors test of normality at the 5% significance level, or Wilcoxon test otherwise; Bonferroni corrected) or in mean preferred phase accordingly (*P* < 0.05; parametric Watson-Williams test for equal means of circular data; Bonferroni corrected). **g**, Same as **f** for pooled NS units of WT and KO mice. NS units from KO animals exhibit, on average, weaker phase-locking, shifted to later phases than those of WT animals, in both delta and theta frequency LFP oscillations.

## DISCUSSION

Multiple studies have shown that a number of mouse models of ASD display various degrees of disrupted excitatory and inhibitory neurotransmission^39,40^ and that a shift in the balance of excitation/inhibition, especially in the medial prefrontal cortex (mPFC), underlie some of the pathologies associated with autism and other psychiatric disorders^25,28^. Here, we find that loss of CNTNAP2, which causes cortical dysplasia focal epilepsy (CDFE), a syndromic form of autism, leads to decreased excitatory and inhibitory synaptic inputs to L2/3 pyramidal neurons in the medial prefrontal cortex (mPFC). LSPS mapping revealed a dramatic reduction of both excitatory and inhibitory inputs in this region. In addition, mEPSCs occurred at a lower frequency in L2/3 neurons, while short-term plasticity and NMDA/AMPA ratios of evoked EPSCs and intrinsic excitability were not altered. These findings suggested a decrease in the number of excitatory synapses, which was confirmed as a decrease in spine density on L2/3 neurons using confocal microscopy, and a decrease in both excitatory and inhibitory synapses using electron microscopy. *In vivo*, these changes were associated with a decreased phase-locking magnitude and shifted phase preference of putative excitatory neurons to delta oscillations and of inhibitory neurons to delta and theta oscillations during locomotion. We conclude that loss of CNTNAP2 has a profound impact on synaptic connectivity and population dynamics of excitatory and inhibitory neurons in the mPFC.

We observed decreased excitatory and inhibitory functional synaptic connectivity and decreased density excitatory and inhibitory synapses in medial prefrontal cortex of Cntnap2 KO animals. Work in dissociated neuronal cultures first showed decreased mEPSC and mIPSC frequency after RNAi knock-down of CNTNAP2, which was associated with decreased dendritic arborization complexity and decreased spine head size^22^. While in general agreement with our findings, showing decreased connectivity, this differs from what we observed in the intact mPFC, as we found no changes in dendritic arborization yet a clear decrease in spine density. In addition, we found no significant changes in spine size or in synapse length in Cntnap2 KO mice, as assessed by electron microscopy L2/3 of mPFC. We also found a significant decrease in MSBs and an increase in perforated PSDs, which further suggests and supports the idea of a complex role of CNTNAP2 in the synaptic cleft. As both MSBs and perforated PSDs reflect well-developed synapses, this change in opposite directions might also indicate the role of both pre-and postsynaptic mechanisms contributing to disruptions in spine maturation dynamics^32,41–44^. These inconsistencies may arise because of differences between cultured neurons and *in vivo* preparations, and may also reflect differences associated with brain regions examined, developmental time-points, or the extent of CNTNAP2 reduction in each experimental condition. Accordingly, recent work in cultured cortical neurons from KO mice does show a decrease in spine density^18^, in agreement with our work. This study also showed decreased localization of the AMPA-subtype glutamate receptor GluA1 in spines of Cntnap2 KO neurons, consistent with the small decrease in mEPSC amplitude we observe in our study. Our results are also in line with our previous work, where we found increased spine elimination and decreased spine density in the apical dendrites of Layer 5 neurons in the somatosensory cortex of Cntnap2 KO mice, using chronic 2-photon imaging *in vivo*^17^. Therefore, the observed effects of CNTNAP2 loss on spine density may not only be restricted to L2/3 of the mPFC and could possibly generalize as decreased spine stability throughout the cerebral cortex.

In contrast, the decrease in inhibitory synapses observed in our study could possibly be related to a reduction in the number inhibitory cells in the Cntnap2 KO cortex, which seems to preferentially impact PV+ neurons, as reported by us and others^20,45^. It has also recently been reported that Cntnap2 KO mice show an age-dependent decrease in spontaneous IPSC frequency and a decrease in tonic GABAergic currents in pyramidal neurons of the visual cortex^46^, all in general agreement with our findings in mPFC. There is also some evidence that decreased synaptic connectivity can also be found outside of cerebral cortex. For example, in CA1, evoked perisomatic inhibition and mIPSC frequency onto CA1 pyramidal neurons is also decreased in Cntnap2 KO^21^.

Interestingly, we found no changes in the intrinsic excitability of L2/3 pyramidal neurons or PV+ neurons in our study. This was surprising at first, given that CNTNAP2 has been shown to be important for axonal potassium channel localization^16^. Brumback et al. found a decrease in input resistance and intrinsic excitability of L5 subcortical projecting neurons^47^, suggesting that laminar location and projection-type of neurons can be an important determinant of altered excitability in neurodevelopmental disease models.

Such alterations in synaptic physiology and neurotransmission seem to be a common theme among mouse models of neurodevelopmental disorders. Shank3, MECP2, and Ube3a loss of function mutations (modeling Phelan McDermid, Rett, and Angelman syndromes, respectively), for example, all result in decreased spine density and decreased excitatory transmission in the cerebral cortex^48–51^, while spine maturation is impaired in the Fragile X model mice^52^, similar to what we find in our Cntnap2 KO. Cortical inhibitory neurotransmission is also similarly compromised in a number of these disorders, as multiple studies have shown^53–56^. This posits the notion that increasing or modulating excitatory and inhibitory synaptic connectivity, especially in a cell-type and projection-specific manner, may be therapeutically relevant.

Concurrently, we find that loss of excitatory and inhibitory synaptic connectivity in Cntnap2 KO mice is associated with a decrease in the magnitude of inhibitory and excitatory neuron phase-locked firing to delta oscillations *in vivo*. Inhibitory neurons were less phase-locked to both delta and theta oscillations and they tended to fire later in the oscillatory cycle. These findings were more prominent during locomotion, which could have broad implications for brain function and suggest that autism relevant-changes in connectivity can be more prominent during specific conditions or arousal states. Temporally-precise firing of different cell types to brain-state specific oscillations has been hypothesized to play a critical role in cortical function^35^. Oscillatory activity in the prefrontal cortex has been extensively studied due to the region’s implications in higher order cognitive functions, including social cognition, working memory, and
attention^24,57^. Moreover, dysfunctional oscillations have often been reported in human individuals diagnosed with ASD and have often been proposed as biomarkers^58–60^. Delta (specifically 4 Hz) oscillations in mPFC can entrain other brain regions such as the amygdala during fear expression, and ventral tegmental area (VTA) and hippocampus during working memory^61^. Theta (4-8 Hz) oscillations in the mPFC, on the other hand, have been associated with signaling safety in conditions of learned fear^62^. Thus, the phase-locking alterations observed in mPFC neurons of Cntnap2 KO mice could be linked to some of the cognitive and affective behavioral disruptions displayed by the model. This posits the potential use for real-time closed-loop neuromodulatory strategies to improve social and cognitive behaviors in autism, warranting further investigation.

The electrophysiological alterations we observe in the mPFC of Cntnap2 KO mice could underlie some of the autism-related phenotypes in the model, such as deficits in social interactions and communication. This is supported by the work of Yizhar et al., who showed that increasing the ratio of excitation to inhibition in the mPFC could in fact disrupt social interactions in WT mice^25^. Moreover, opsin-mediated increase in PV cell excitability, or decrease in pyramidal neurons of prelimbic mPFC has already been shown to successfully rescue social behavior and hyperactivity in Cntnap2 KO mice^28^. Such disruptions in excitatory/inhibitory balance could also be reflected in broader scale as alterations in oscillatory power and synchrony. Moreover, these synaptic alterations could be mechanistically linked to the altered representation of social stimuli in the mPFC of Cntnap2 KO mice^63^. Consistent with this notion, a number of syndromic forms of autism show disrupted oscillations, such as Angelman Syndrome, which presents with enhanced delta oscillations^58^, and Duplication 15q Syndrome, which shows enhanced beta oscillations^64^.

Future studies will need to dissect the inputs and outputs of the prefrontal cortex in a cell-type and projection-specific manner, in order to discover whether changes in excitatory and inhibitory connectivity are generalized or selectively impaired in specific circuits. Also, it is not known whether the synaptic and population dynamic changes we have found can be reversed or ameliorated by restoring CNTNAP2 gene expression in adulthood, or whether very early interventions will be needed. Finally, it will be important to understand how the delta and theta phase-locking impacts recruitment of other connected brain regions, especially in the context of social engagement.

## METHODS

**Animals.** Cntnap2 null mice were obtained from E. Peles and backcrossed to the C57BL/6J background for over 12 generations. For targeted electrophysiological recordings of parvalbumin-positive interneurons, Cntnap2 heterozygous mice were backcrossed to PV-Cre (Jackson labs number 008069) x Ai9 (Jackson labs number 007909) mice. For spine density analysis, Cntnap2 heterozygous mice were backcrossed with Thy1GFP (Jackson labs x 007788) mice. Experimental mice were obtained from heterozygous crossings and born with the expected Mendelian frequencies; both genders were used. The date of birth was designated at P0 and the three obtained genotypes (wild-type, heterozygous, homozygous knock-out) were housed together with three to four mice per same-sex cage. Mice were kept in a 12-hour light/12-hour dark cycle and had ad libitum access to food and water. All procedures involving animals were performed in accordance with the University of California, Los Angeles (UCLA) animal research committee, and the National Institutes of Health *Guide for the Use and Care of Laboratory Animals*.

**Slice preparation.** Acute coronal slices (300 μm thickness) containing the medial prefrontal cortex were prepared from 4 to 6-wk-old Cntnap2 knock-out mice and wild-type littermates. Mice were anaesthetized with isoflurane gas and beheaded after disappearance of toe-pinch reflex. The brain was removed and placed in ice-cold cutting solution consisting of (mM): 222 sucrose, 11 D-glucose, 26 NaHCO_3_, 1 NaH_2_PO_4_, 3 KCl, 7 MgCl_2_, 0.5 CaCl_2_, aerated with 95% O_2_, 5% CO_2_. The brain was cut in a Leica VT1000S Vibratome. Slices were allowed to recover for 30 minutes at 37 °C in standard artificial cerebrospinal fluid (ACSF, in mM): 124 NaCl, 2.5 KCl, 26 NaHCO_3_, 1.25 NaH_2_PO_4_, 10 D-glucose, 4 sucrose, 2.5 CaCl_2_, 2 MgCl_2_, aerated with 95% O_2_, 5% CO_2_, and kept at room temperature for at least 40 min until time of recording.

**Electrophysiology.** Whole-cell patch-clamp recordings of L2/3 neurons were obtained under visual guidance using infrared DIC video-microscopy and water-immersion 40x objective, with patch pipettes (3-5 MOhms) pulled from borosilicate capillary glass (Sutter) with a Sutter puller. Tdtomato-expressing parvalbumin-positive inhibitory neurons were targeted under epifluorescence. All electrophysiological recordings were performed using Multiclamp 700B (Molecular Devices) patch clamp amplifiers and ACSF was maintained at 33–35 °C. Signals were filtered at 4 kHz using Bessel filter and digitized at 10 kHz with WinWCP and WinEDR electrophysiology software interface for voltage-clamp recordings (Strathclyde). Current clamp recordings were digitized at 15 and Bessel filtered at 6 kHz. Series/access resistance was monitored in all recordings and compensated in current clamp mode. Recordings were discarded if series resistance changed significantly (>20%) or exceeded 25 MOhms. Junction potential was not compensated.

**Current-clamp recordings.** For intrinsic excitability experiments, the internal pipette solution contained (in mM): 115 KGluc, 20 KCl, 10 HEPES, 10 phosphocreatine, 4 ATP–Mg^2+^, 0.3 GTP–Na^+^ (pH 7.2, 270-290 mOsm); in some recordings, 0.2% biocytin was added to the solution. Patched pyramidal excitatory neurons were identified and included in the analysis based on their action potential firing characteristics. Resting membrane potential (V_m_) was measured after breaking into the cell (rupturing the patch) and applying zero current. Input resistance (R_in_) was calculated as the slope of the linear fit of the voltage-current plot, generated from a family of negative and positive 500 ms current injections (−60 pA to +60 pA at 20 pA intervals, for pyramidal cells; −150 pA to +150 pA at 50 pA intervals, for parvalbumin-positive interneurons). The membrane decay constant (τ) was calculated by fitting a single exponential curve to the current-voltage plot that resulted from a −20 pA current injection. Cell membrane capacitance (C_m_) was given by C_m_ = τ/R_in_. For assessment of intrinsic excitability, cells were clamped at −70 mV and injected a series of increasing current steps at 50 pA intervals. Action potential properties were determined from the first action potential elicited by minimum current injection. The spike adaptation ratio was calculated by dividing the last inter-spike interval to the first inter-spike interval in an action potential train elicited by a 500 ms pulse of 200 pA. All data was analyzed using custom-written MATLAB software.

**Voltage-clamp recordings.** Miniature excitatory postsynaptic currents (mEPSCs) were isolated by applying (in mM): 0.5 tetrodotoxin (TTX) and 10 pictrotoxin to ACSF (described above). Pipette internal solution contained (in mM): 20 KCl, 10 Na-phosphocreatinine, 100 cesium methyl sulfonate, 3 QX-314, 10 HEPES, 4 ATP–Mg^2+^ and 0.3 GTP–Na^+^ (pH 7.2, 270-290 mOsm). Recordings were performed with cells clamped at −70 mV. Miniature inhibitory postsynaptic currents (mIPSCs) were isolated by applying (in mM): 0.5 tetrodotoxin (TTX), 10 CNQX, and 50 APV to ACSF. A high-chloride pipette internal solution was used, which contained (in mM): 120 KCl, 10 HEPES, 4 ATP-Mg^2+^, 0.3 GTP–Na^+^ and 10 Na-phosphocreatinine (pH 7.2, 270-290 mOsm). Recordings were performed with cells clamped at −50 mV. Miniature and spontaneous events were recorded for 2 min. MiniAnalysis software (Synaptosoft) was used to automatically identify synaptic events, based on template parameters. Events were then manually examined to exclude false positives. For voltage clamp recordings with a cesium-containing electrode, pyramidal cells were targeted based on soma shape and identity was manually verified based on EPSC decay, where cells with mean ESPC decay time constant ≤2 ms were considered to likely inhibitory and excluded. Events were excluded if the 10-90% rise time was > 2 ms, as these events were likely recorded from synapses far from the soma and with poor space clamp. Inter-event intervals (event frequency), amplitude, decay time constant, area, 10-90% rise time, and half-width, were analyzed and comparisons between groups were analyzed by Student’s T-test. Grouped data are expressed as mean ± SEM.

**Evoked Excitatory Postsynaptic Currents.** A tungsten concentric bipolar stimulating electrode (WPI) was placed in the white matter to stimulate axon fibers emerging from the anterior forceps of the corpus callosum, which project onto a whole-cell recorded L2/3 pyramidal neuron in PL-mPFC, voltage-clamped at −70 mV. Input-output curves were derived by increasing the stimulus duration (0.1 ms increments) and recording current responses in the recorded postsynaptic neurons. Short-term plasticity was assessed by measuring paired-pulse ratios, calculated as the peak amplitudes of 10 averaged episodes at various inter-stimulus intervals (25, 50, 100, 500ms). AMPA/NMDA ratios were measured by voltage-clamping the cells at a holding potential of −70 mV for AMPA currents and +40 mV for NMDA currents. Peak amplitude current responses were averaged over 10 episodes. Peak NMDA currents were measured after the offset of AMPA currents (25-30 ms post-stimulus) within the same cell. Data was analyzed manually using WinEDR software and plotted in MATLAB.

**Laser Scanning Photostimulation (LSPS).** Coronal sections of medial prefrontal cortex were cut 400 μm thick with a vibratome (VT1200S, Leica Systems) in sucrose-containing artificial cerebrospinal fluid (ACSF) (in mM: 85 NaCl, 75 sucrose, 2.5 KCl, 25 glucose, 1.25 NaH2PO4, 4 MgCl2, 0.5 CaCl2, and 24 NaHCO3). Slices were first incubated in sucrose-containing ACSF for 30 min to 1 h at 32°C, and then transferred to recording ACSF (in mM: 126 NaCl, 2.5 KCl, 26 NaHCO3, 2 CaCl2, 2 MgCl2, 1.25 NaH2PO4, and 10 glucose) at room temperature. Throughout incubation and recording, the slices were continuously bubbled with 95% O2-5% CO2. The design of our laser scanning photostimulation system has been described previously^65^. A laser unit (model 3501, DPSS Lasers, Santa Clara, CA) was used to generate a 355 nm UV laser for glutamate uncaging. Various laser stimulation positions were achieved through galvanometer-driven X-Y scanning mirrors (Cambridge Technology, Cambridge, MA), as the mirrors and the back aperture of the objective were in conjugate planes, thereby translating mirror positions into different scanning locations at the objective lens focal plane. Data were acquired with a Multiclamp 700B amplifier (Molecular Devices, Sunnyvale, CA), data acquisition boards (models PCI MIO 16E-4 and 6713, National Instruments, Austin, TX), and custom-modified version of Ephus software (Ephus, available at https://www.ephus.org/). Data were low-pass filtered at 2 kHz using a Bessel filter, digitized at 10 kHz, and stored on a computer. Cortical slices were visualized with an upright microscope (BW51X, Olympus) with infrared differential interference contrast optics. Electrophysiological recordings, photostimulation, and imaging of the slice preparations were done in a slice perfusion chamber mounted on a motorized stage of the microscope at room temperature. An aliquot of MNI-caged-L-glutamate (4-methoxy-7-nitroindolinyl-caged L-glutamate, Tocris Bioscience, Ellisville, MO) was added to 20–25 mL of circulating ACSF for a concentration of 0.2 mM caged glutamate. To perform whole cell recording, cells were visualized at high magnification (60× objective, 0.9 NA; LUMPlanFl/IR, Olympus). Excitatory neurons were selected based upon their pyramidal somata detected under differential interference contrast (DIC) microscopy. For experiments to assess photo-stimulation evoked spiking profiles of excitatory in mPFC (similar to our published studies^65,66^), the patch pipettes (4–6 MΩ resistance) were filled with an K+ internal solution containing (in mM) 126 K-gluconate, 4 KCl, 10 HEPES, 4 ATP-Mg, 0.3 GTP-Na, and 10 phosphocreatine (pH 7.2, 300 mOsm). For the photostimulation experiments to map synaptic inputs, we used a Cs+ internal solution containing (in mM) 6 CsCl, 130 CsOH, 130 D-Gluconic acid, 2 MgCl2, 0.2 EGTA, 10 HEPES, 2.5 ATP-Na, 0.5 GTP-Na, and 10 phosphocreatine-Na2 (pH 7.2, 300 mOsm). Because glutamate uncaging agnostically activates both excitatory and inhibitory neurons, we empirically determined the excitatory and inhibitory reversal potentials in L2/3 pyramidal cells to properly isolate EPSCs and IPSCs. Whole-cell voltage-clamp recordings were made from the recorded postsynaptic neurons with LSPS-evoked EPSCs and IPSCs measured at the holding potential of −70 mV and +5 mV, respectively, across photostimulation sites. The internal solution also contained 0.1% biocytin for cell labeling and morphological identification. The morphology of recorded pyramidal neuron was determined using post-hoc staining with Cy3-conjugated streptavidin (1:500 dilution; Jackson ImmunoResearch). Once stable whole cell recordings were achieved with good access resistance (usually <30 MΩ), the microscope objective was switched from 60× to 4×; laser scanning photostimulation (LSPS) was performed through the 4x objective lens. At low magnification (4× objective lens, 0.16 NA; UplanApo, Olympus), the slice images were acquired by a high-resolution digital CCD camera (Retiga 2000, Q-imaging, Austin, TX) and used for guiding and registering photostimulation sites in cortical slices.

Photostimulation (1.5 ms duration, 15 mW pulses) from a 350 nm UV laser generator (DPSS Lasers, Santa Clara, CA) was delivered to the sample, controlled via an electro-optical modulator and a mechanical shutter. Focal laser spots approximated a Gaussian profile with a diameter of ~50-100 μm. Under our experimental conditions, LSPS evoked action potentials were recorded from stimulation locations within 100 μm of targeted somata of excitatory neurons and occurred within 150 ms post photostimulation. Our calibration analysis indicates that LSPS allows for mapping direct synaptic inputs to recorded neurons. Synaptic currents in patched neurons were detected under voltage clamp. By systematically surveying synaptic inputs from hundreds of different sites across a large cortical region, aggregate synaptic input maps were generated for individual neurons. For our mapping experiments, a standard stimulus grid (16×16 stimulation sites, 100 × 60 μm^2^ spacing) was used to tessellate mPFC from pia to white matter. The LSPS site spacing was empirically determined to capture the smallest predicted distance in which photostimulation differentially activates adjacent neurons. Glutamate uncaging was delivered sequentially in a nonraster, nonrandom sequence, following a “shifting-X” pattern designed to avoid revisiting the vicinity of recently stimulated sites.

Photostimulation induces two forms of excitatory responses: (1) those that result from direct activation of the recorded neuron’s glutamate receptors, and (2) synaptically mediated responses (EPSCs) resulting from the suprathreshold activation of presynaptic excitatory neurons. Responses that occur within 10 ms of laser pulse onset were considered direct; these responses exhibited a distinct waveform and occurred immediately after glutamate uncaging. Synaptic currents with such short latencies are not possible because they would have to occur before the generation of action potentials in photostimulated neurons. Therefore, direct responses were excluded from local synaptic input analysis, but they were used to assess glutamate mediated excitability/responsiveness of recorded neurons. At some locations, synaptic responses were overriding on the relatively small direct responses, and these responses were identified and included in synaptic input analysis. The IPSC input was similarly analyzed as the EPSC input. For data map analysis, we implemented the approach for detection and extraction of photostimulation-evoked postsynaptic current responses as previously described^66^. LSPS evoked EPSCs/IPSCs were quantified across the 16x16 mapping grid for each cell, and 1-2 individual maps were used per recorded cell. The PSC input from each stimulation site was the measurement of the sum of individual PSCs within the analysis window (>10 ms to 160 ms post photostimulation), with the baseline spontaneous response subtracted from the photostimulation response of the same site. The value was normalized with the duration of the analysis window (i.e., 150 ms) and expressed as average integrated amplitudes in picoamperes (pA). The analysis window was chosen because photostimulated neurons fire most of their action potentials during this time. For the color-coded map display, data were plotted as the average integrated PSCs amplitude per pixel location (stimulation site), with the color scale coding input strength. For the group maps obtained across multiple cells, the individual cell maps were first aligned by their slice images using laminar cytoarchitectonic landmarks. Then a new map grid was created to re-sample and average input strength at each site location across cell maps; a smooth version of color-coded map was presented for overall assessments. To further quantitatively compare input strength across cell groups, we measured the total PSC inputs (total synaptic currents) across all map sites (total synaptic input strength) for individual cells. The total EPSC/IPSC input strength ratios were also measured for the cells when both EPSC and IPSC data were available from the same cells.

As virtually all Layer 1 neurons are inhibitory cells, and pyramidal neurons with apical dendritic tufts in layer 1 could fire action potentials when their tufts were stimulated in layer 1^67^, EPSCs detected after photostimulation in layer 1 were not included for analyses. However, because layer 1 neurons can provide inhibition to layer 2/3 neurons, we did analyze IPSCs detected after photostimulation in layer 1. All data are reported as mean ± standard error of the mean (SEM). When comparing two independent groups, a Mann-Whitney U test was used. Unless specified otherwise, sample size n was defined as cell number. A *P* value (≤ 0.05) was considered statistically significant.

**Immunohistochemistry.** For assessment of dendritic morphology and complexity, cells were during electrophysiological recordings via passive diffusion of internal pipette solution-containing 0.2% biocytin. After recording for at least 10 min, slices were transferred to a 4% PFA solution for overnight fixation, washed for 10 min (x3) in 0.1 M phosphate buffered saline (PBS), blocked with 10% normal goat serum (NGS) containing 0.3% Triton-X in 0.1 M PBS for 1.5 hrs, and incubated overnight with an Alexa 555 or Alexa 488-conjugated Streptavidin antibody (1:500, Invitrogen) in 0.1M PBS. Sections were finally washed 3x 10 min in 0.1M PBS and mounted on slides using DAPI Fluoromount-G (Invitrogen) for visualization. We assessed dendritic complexity of biocytin-filled cells by imaging at 20X magnification in an LSM 520 confocal microscope. Z-stacks of optical sections (1 um) were compiled and images were processed in Neurolucida 10 (MFB Biosciences) for Sholl analysis.

For quantification of spine density, Cntnap2 WT and KO mice were crossed with a Thy1-GFP mouse line, which sparsely labels pyramidal neurons, including their dendritic projections and spines. Mice were perfused intracardially with 25 mL 0.1 M PBS, followed by 25 mL of 4% PFA in 0.1 M PBS (at 2 mL/min). The brains were dissected and fixed for at least 24 hrs in the same solution. Brains were then sectioned at a thickness of 100 um, using a Leica vibratome. Sections containing the mPFC were mounted in slides using DAPI Fluoromount-G media. Apical and basal dendrites of GFP-expressing L2/3 mPFC neurons were imaged at high resolution using a 63X oil magnification objective on an LSM 520 confocal microscope (Zeiss). Optical sections of 0.32 um were acquired and maximum intensity projections of dendritic arbors were created in ImageJ. Dendritic segments were chosen using consistent criteria and spines were manually counted. Dendritic density was calculated by dividing the total number of spines over a given length of dendrite (spines/μm). Student’s t-test was performed for statistical comparison between WT and KO mice.

**Tissue preparation and electron microscopy.** Animals from KO and WT groups (n = 3, respectively) were processed. Mice were deeply anesthetized with isoflurane were perfused transcardially with a mixture of 2% paraformaldehyde (PFA) and 2% glutaraldehyde in 0.1 M phosphate buffer (PB, pH 7.4). Brains were removed and post-fixed overnight at 4°C. 70 micrometer thick sections were cut with a Leica vibratome. Free-floating sections for electron microscopy were post-fixed with 1% OsO_4_, dehydrated in ascending ethanol series and embedded in epoxy resin (Durcupan; Sigma, Germany) within Aclar sheets (EMS, Hatfield, PA, USA). Uniform rectangular samples were cut from the prelimbic medial prefrontal cortices (mPFC, approx 1.98-1.94 AP, 3-3.75 DV, 0.25-0.50 ML position) under a Leica S6E dissecting microscope, and mounted on plastic blocks. 60 nm ultrathin sections were cut on a Reichert ultramicrotome, mounted on 300 mesh copper grids, contrasted with lead citrate (Ultrostain II, Leica) and examined with a JEM-1011 transmission electron microscope (JEOL, Tokyo, Japan) equipped with a Mega-View-III digital camera and a Soft Imaging System (SIS, Münster, Germany) for the acquisition of the electron micrographs. Five to ten sections were analyzed per block, and two blocks per animal were used to collect micrographs. Sample areas (at least 50 μm^2^ per animal) were chosen in a pseudo-random fashion and photographed at a uniform magnification. Postsynaptic dendritic spines, axonal boutons, multi-synaptic boutons (MSB; a single presynaptic bouton that forms separate synapses with multiple spine heads) were identified on electron micrographs. Spine profile area were measured using the engine provided by NIH ImageJ v1.51j8 (Schneider et al., 2012); data were compiled using Excel (Microsoft) and Kaleidagraph (Synergy Software, Reading, PA, USA) software. The means and the effects of the loss of the Cntnap2 gene was determined by Mann-Whitney. U test, with a P < 0.05 considered statistically significant. Data collection and quantification was performed blindly, to eliminate bias. We performed electron microscopy using random sampling from single sections to optimize sample size and to detect changes in synaptic features associated with loss of Cntnap2 in KO mice, compared to WT.

**Surgery, behavioral habituation, and *in vivo* electrophysiology.** Adult male and female Cntnap2 mutant and wild-type mice (2-5 months old) underwent an initial surgery for implantation of a stainless steel head restraint bar on their skull in preparation for *in vivo* electrophysiological recordings. All surgical procedures were performed under isoflurane anesthesia (3–5% induction, 1.5% maintenance) in a stereotaxic apparatus. Mouse body temperature was monitored and kept at 37°C during surgery using a Harvard Apparatus feedback-controlled heating pad and were administered an subcutaneous injection of carprofen (5 mg/kg of body weight) for systemic analgesia. Mice were allowed to recover for 5 days, during which they were given antibiotic treatment (amoxicillin, 0.25 mg/mL in drinking water). After recovery period, mice were habituated for at least three days for each of the following stages: human handling (5 min), headbar attachment (10 min), and head fixation on a spherical treadmill (10 min). The treadmill consisted of an 8-inch Styrofoam ball (Graham Sweet), tethered with a metal rod through the middle allowing only one axis of rotation. Air was blown, allowing the ball to float and the mouse to spin the ball and run in place and on top of it (Polack et al, 2013). After habituation, and one day prior to electrophysiological recordings, the mouse received a craniotomy above the medial prefrontal cortex on the right hemisphere (anterior 1.8 mm, lateral 0.5 mm to Bregma). The dura above the exposed brain area was carefully removed in order to facilitate electrode insertion. The exposed skull and brain were covered and sealed with a silicone elastomer sealant (Kwik-Sil, WPI). An additional craniotomy was performed over the posterior cerebellum for placement of a silver chloride electrical reference wire, which was glued into place with dental cement. The mouse was allowed to recover overnight. Mice were given a dose of carprofen on day of recording, to ameliorate any pain associated with the craniotomy surgery.

On the day of the recording, the mouse was head-fixed atop the spherical treadmill, the Kwik-Sil was removed and cortex buffer (135 mM NaCl, 5 mM KCl, 5 mM HEPES, 1.8 mM CaCl_2_ and 1 mM MgCl_2_) was immediately placed on top of the craniotomy in order to keep the exposed brain moist. The mouse skull was then stereotaxically aligned and the silicon microprobe coated with a fluorescent dye (DiI, Invitrogen), was stereotaxically lowered using a micromanipulator into the mPFC (relative to bregma: anterior 1.8 mm, lateral 0.5 mm, ventral 2.5 mm). This process was monitored using a surgical microscope (Zeiss STEMI 2000). The microprobes contained a total of 128 electrode recording sites that were densely distributed (hexagonal array geometry with 25 mm vertical spacing and 16-20 mm horizontal spacing) on two prongs (placed 0.4 mm apart), spanning L2/3 and L5 of the prelimbic (PL) and infralimbic (IL) medial prefrontal cortex. Only data from L2/3 prelimbic cortex was used. Once inserted, the probe was allowed to settle among the brain tissue for 1 hr. Recording of brain network activity was done for a total duration of 1 hr after that.

Data acquisition was performed using custom fabricated silicon probes and recorded with LabView Software^68^. Readout was achieved via a custom-built 128-channel detachable head stage module. Head stages contained commercial integrated electronic circuits (Intan Technologies RHA-2164B)^69^ providing signal multiplexing (32 electrodes per multiplexed output wire), amplification (gain 200), and filtering (0.1– 6500 Hz) functions. The head stage contained two 64-pin connectors (Molex, Slimstack 502426-6410) connecting to custom printed circuit boards wire bonded to the silicon microprobes. Analog signals were transmitted through thin flexible cables and subsequently digitized on 16-bit analog-to-digital conversion (ADC) cards (USB-6356, National Instruments). Multiplexed signals were recorded at 800 kHz and de-multiplexed with recording software into a sampling rate of 25 kHz per channel. All ADC cards were synchronized via a shared internal clock. All data acquisition, as well as control of stimulus timing, was performed with custom LabVIEW scripts. All data analysis was carried out with custom MATLAB scripts^36^.

After the recording session, mice were anaesthetized with isoflurane and sacrificed. The brain was extracted, sectioned (100 μm) on a Vibratome (Leica) and mounted on slides with DAPI Fluoromount-G (SouthernBiotech) mounting media. Confocal tiled images were taken to verify microprobe location (Zeiss LSM 800). Anatomical landmarks were used to determine anterior-posterior coordinates relative to bregma. Each of the 128 recording sites was then assigned an approximate coordinate in 3D Cartesian space and classified as belonging to prelimbic (PL) or infralimbic (IL) prefrontal cortex (Allen Brain Atlas).

**Motion detection.** Mouse treadmill rotation was recorded as an analog signal, using a custom printed circuit board based on a high sensitivity gaming mouse sensor (Avago ADNS-9500) connected to a microcontroller (Atmel Atmega328). The signal was initially recorded along with electrophysiology at 25 kHz, then downsampled to 1 kHz, its sample mode value was subtracted, it was turned to absolute values and was smoothed by lowpass filtering <1 Hz with a first order Butterworth filter. For one set of animals (n = 6) motion was detected when the smoothed treadmill motion-signal exceeded 0.8 × mean of recording and immobility was assumed when it dropped below 0.005 (a.u). For a second set (n = 7) these thresholds were changed to 2 x mean and 0.016 respectively due to increased recording noise. Motion segments shorter than 0.5 s long were discarded, and consecutive segments closer than 0.5 s were concatenated.

**Local Field Potential Analysis.** To compare LFP bandpower, LFP recordings from the channel located closest to L2/3 were selected for each animal. LFPs were down-sampled to 1 kHz and all data points corresponding to either motion or immobility segments were concatenated. Bandpower over all frequency ranges (delta: 1-4 Hz, theta: 5-8 Hz, beta: 12-30 Hz, slow gamma: 30-55 Hz, fast gamma: 80-110 Hz) was computed using a periodogram with a Hamming window.

***In vivo* Unit Clustering and Analysis.** Single units were isolated using custom spike-detection scripts and PyClust software^68^. Raw data was initially background subtracted, bandpass filtered from 600-6500 Hz, and grouped into channel sets of neighboring electrodes for clustering. For each channel set, putative spikes were detected as any deviation greater than 4 standard deviations from the mean. The features of each spike were calculated (peak amplitude, valley, and trough, principle components) and individual clusters were isolated by outlining boundaries on each projection. Each unit was visually inspected and units that drifted outside recorded channels or were lost during the recording were eliminated. For each clustered unit, all peri-spike extracellular waveforms were collected from the channel that yielded the largest spike amplitude and were bandpass filtered at 600 - 6000 Hz with a first order Butterworth filter. The amplitudes and time-points of each unit’s mean waveform peak and trough were computed, together with the unit’s mean firing rate and burst index (percentage of consecutive spikes closer than 20 ms). Waveforms from identified units were visually inspected and units with low-quality individual and average waveforms were not included in the analysis (n = 25 rejected units).

To cluster units into broad and narrow spiking, the (i) peak-trough time distance, (ii) peak-trough amplitude ratio and (iii) burst index of all units of all animals were pooled together and z-scored. Principal component analysis was performed on the three variables and the scores from all three principal components were split into two clusters using k-means with squared Euclidean distance measure and 100 clustering repeats. This method yielded two well separated clusters with one containing ~20% of all units with smaller peak-trough distances and ratios and higher burst indexes compared to the other cluster. Units in that cluster are referred to as ‘narrow-spiking’ whereas those in the opposite cluster as ‘wide-spiking’.

The phase of each spike was computed using the LFP recording of the channel yielding the largest spike amplitude for that unit. LFPs were down-sampled at 1 kHz and bandpass filtered over the corresponding oscillation frequency range using a Butterworth bandpass filter matching the passband exactly. The phase of each spike was computed as the angle of the filtered LFP’s Hilbert transform at the spike peak. For producing preferred phase distributions of wide or narrow spiking units, we only included units with > 200 spikes over all motion or immobility segments accordingly and with significant phase locking at the corresponding frequency range (Rayleigh test, p-value < 0.05, Bonferroni corrected over all wide or narrow spiking units accordingly). Moreover, a minority of units in the narrow-spiking cluster with large peak-trough distances (>0.4 ms) falling within the broad spiking cluster, were removed without affecting any results in the analysis.

## ACKNOWLEDGEMENTS

PG, JT, and ML were supported by NIH grants 1R01MH101198, U01 NS094286, R01 MH105427, U54 HD87101. ML was funded by the UCLA Eugene Cota-Robles Fellowship, NSF-GRFP DGE-0707424, the NIMH T32MH073526 UCLA Neurobehavioral Genetics Training Grant, the UCLA Dissertation Year Fellowship, and NIH grant 5U01NS094286. TS was supported by a postdoctoral fellowship from the Epilepsy Foundation. SCM was supported by NIH DA034178. We thank Tünde Magyar and Renáta Pop for excellent technical assistance. The Project is supported by the European Union and co-financed by the European Social Fund (grant agreement no. EFOP-3.6.2-16-2017-0008: The role of neuro-inflammation in neurodegeneration: form molecules to clinics). BR was also supported by the János Bolyai Research Fellowship of the Hungarian Academy of Sciences. x

## AUTHOR CONTRIBUTIONS

M.T.L., D.H.G., and P.G. designed the experiments. M.T.L. carried out slice recordings, neuronal anatomy experiments, *in vivo* experiments, and analyzed data. J.T. analyzed *in vivo* recordings. T.S. helped with experiment design and built the *in vivo* recording setup. I.B. performed surgeries, *in vivo* recordings, and immunohistochemistry. T.I. and R.S. performed the LSPS experiments and analysis, and X.X. helped with experimental design. M.M. and B.R. performed electron microscopy and analysis. A.M. and S.C. performed immunohistochemistry. A.F., R.G., N.S., and D.T. performed spike-sorting in PyClust and helped with *in vivo* recordings. M.A. performed dendrite tracing. H.D. helped with mouse breeding and genotyping. O.P. contributed to experimental design. S.M. provided custom-built silicon microprobes and helped with setting up the *in vivo* electrophysiological recordings.

---

**Supplementary Fig. 1.**
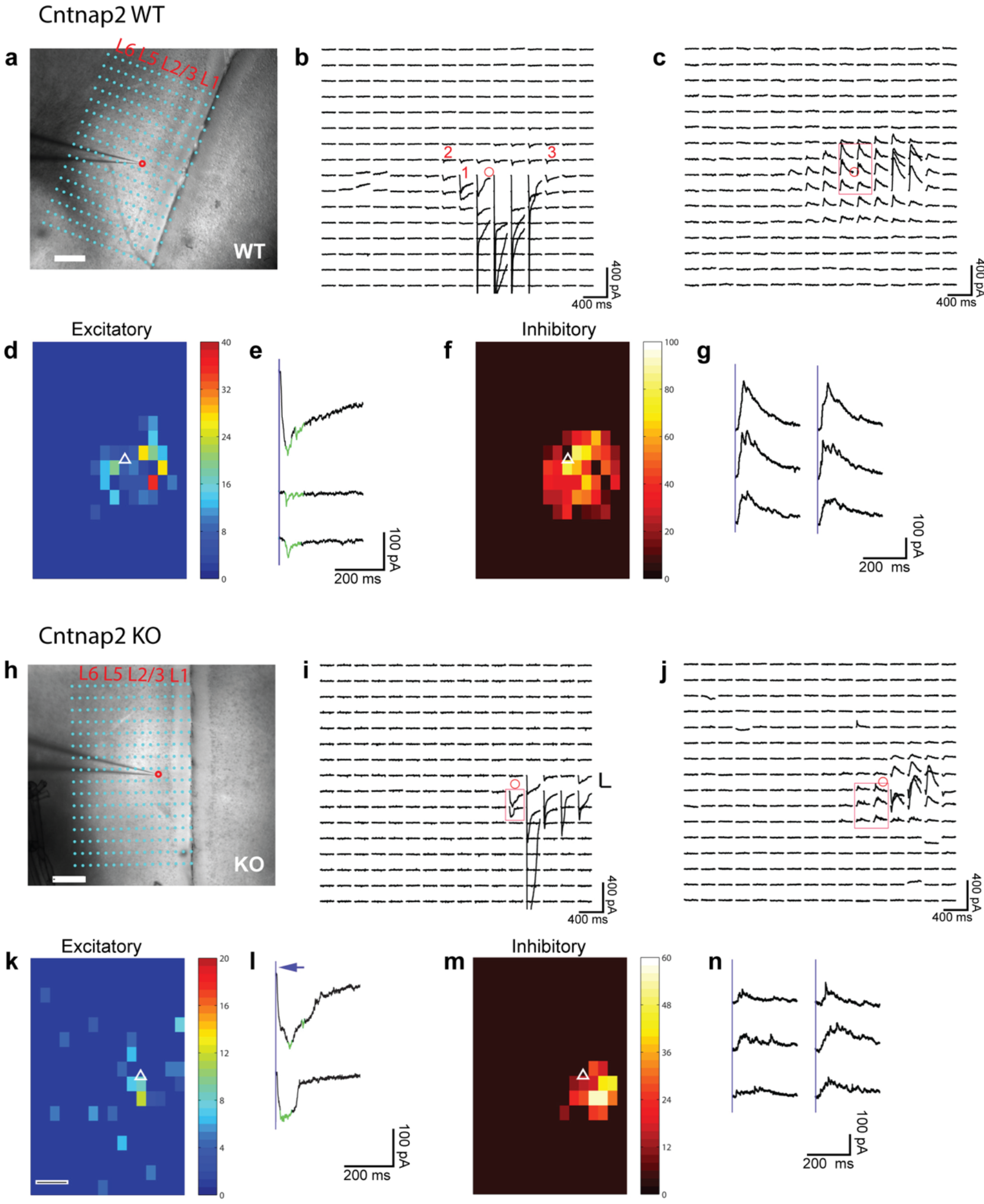
Example cortical input map data for Cntnap2 WT and KO L2/3 mPFC excitatory neurons. **a**,**h**, Differential interference contrast (DIC) image of mPFC, superimposed with photostimulation sites (cyan dots), spaced at 100 μm × 60 μm, for WT and KO mice. The tip of the patch pipette (recording electrode) and the cell body location of a recorded L2/3 neuron is indicated by a red circle. **b**,**c**,**i**,**j**, Photostimulation-evoked response traces plotted according to their corresponding photostimulation sites, as shown in **a**, **h**. Traces depict currents recorded 250 ms after stimulation (1.5 ms, 15 mW) onset. Cells were voltage ‑ clamped at −70 mV to detect inward excitatory postsynaptic currents (EPSCs), depicted in **b** and **i**, and at +5 mV to detect inhibitory postsynaptic currents (IPSCs), depicted in **c** and **j**. Excitatory **d**,**k** and inhibitory **f**, **m** input maps of average integrated stimulation responses for datasets shown in **b**,**i** and **c**,**j**, respectively. Somatic location of the recorded neuron is represented by a white triangle. **e**,**l** and **g**,**n** show enlarged insets of selected responses in **b**,**i** and **c**,**j**, respectively. Green overlays mark over-riding synaptic responses. Average input amplitudes were calculated as mean integrated amplitudes of EPSCs or IPSCs elicited within the 250 ms post-stimulus onset time-frame. White scale bars represent 250 μm.

**Supplementary Fig. 2.**
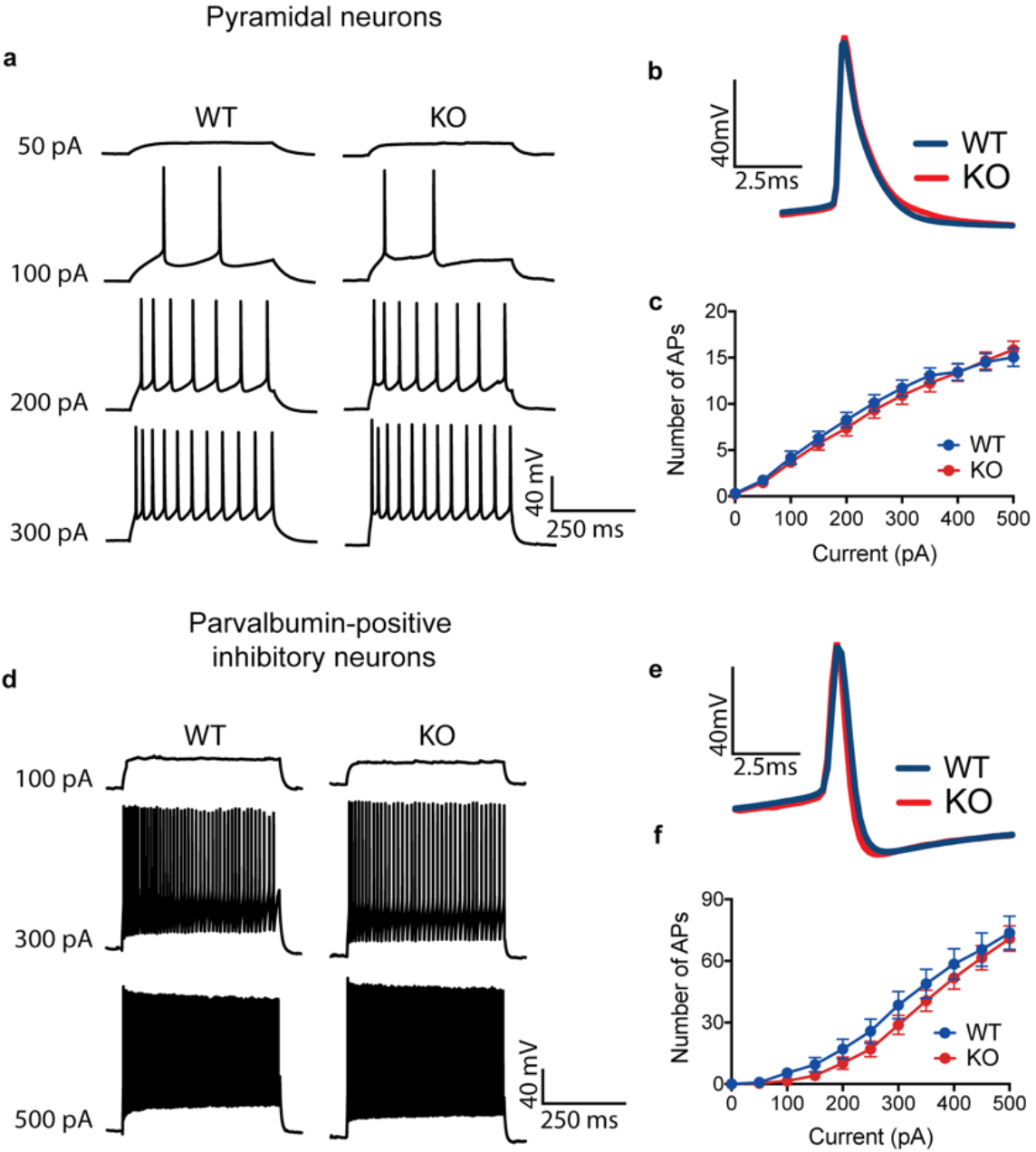
Intrinsic excitability of L2/3 pyramidal neurons and parvalbumin-positive (PV+) inhibitory neurons in Cntnap2 WT and KO mice. **A**,**d,** Representative action potential traces from L2/3 WT and KO pyramidal and PV+ neurons, showing responses to various current injections and **b**,**e**, corresponding average action potential waveforms. **c**,**f**, Input-output curves showing average number of action potentials elicited by increasing current injections for pyramidal neurons (WT *n* = 28 cells, KO *n* = 21 cells; *P* = 0.7057, 2way ANOVA) and PV+ inhibitory neurons (WT *n* = 27 cells, KO *n* = 42 cells; *P* = 0.2993). Data obtained from current-clamp recordings of neuronal spikes elicited by stimulating with 50 pA step increments, cells clamped at −70mV.

**Supplementary Table 1.**

Passive Membrane Properties
| Parameters | Pyramidal Neurons |  | PV Inhibitory Neurons |  |
| --- | --- | --- | --- | --- |
|  | WT | KO | WT | KO |
| RMP (mV) | -73.2 ± 2.0 | -68.7 ± 2.3 | -80.6 ± 1.2 | -78.0 ± 0.8 |
| Rin (mOhms) | 174.1 ± 18.7 | 173.5 ± 21.7 | 93.2 ± 6.3 | 85.1 ± 4.2 |
| Cm (pF) | 96.1 ± 8.1 | 106.8 ± 12.3 | 69.8 ± 4.2 | 80.5 ± 4.5 |
| Tau (ms) | 14.7 ± 1.0 | 16.6 ± 0.9 | 6.2 ± 0.2 | 6.4 ± 0.2 |

Action Potential Features
| Parameters | Pyramidal Neurons |  | PV Inhibitory Neurons |  |
| --- | --- | --- | --- | --- |
|  | WT | KO | WT | KO |
| Amplitude (mV) | 84.9 ± 1.7 | 82.1 ± 1.9 | 62.0 ± 2.3 | 64.5 ± 1.7 |
| Half-width (ms) | 1.0 ± 0.1 | 1.1 ± 0.1 | 0.3 ± 0.0 | 0.3 ± 0.0 |
| AHP Amplitude (mV) | -4.5 ± 1.2 | -4.0 ± 1.7 | -24.0 ± 0.7 | -22.1 ± 0.6 |
| Peak to AHP (ms) | 3.8 ± 0.5 | 3.1 ± 0.5 | 0.8 ± 0.1 | 0.8 ± 0.0 |
| Threshold (mV) | -39.3 ± 1.0 | -37.0 ± 0.9 | -39.7 ± 1.0 | -41.0 ± 0.9 |

**Supplementary Fig. 3.**
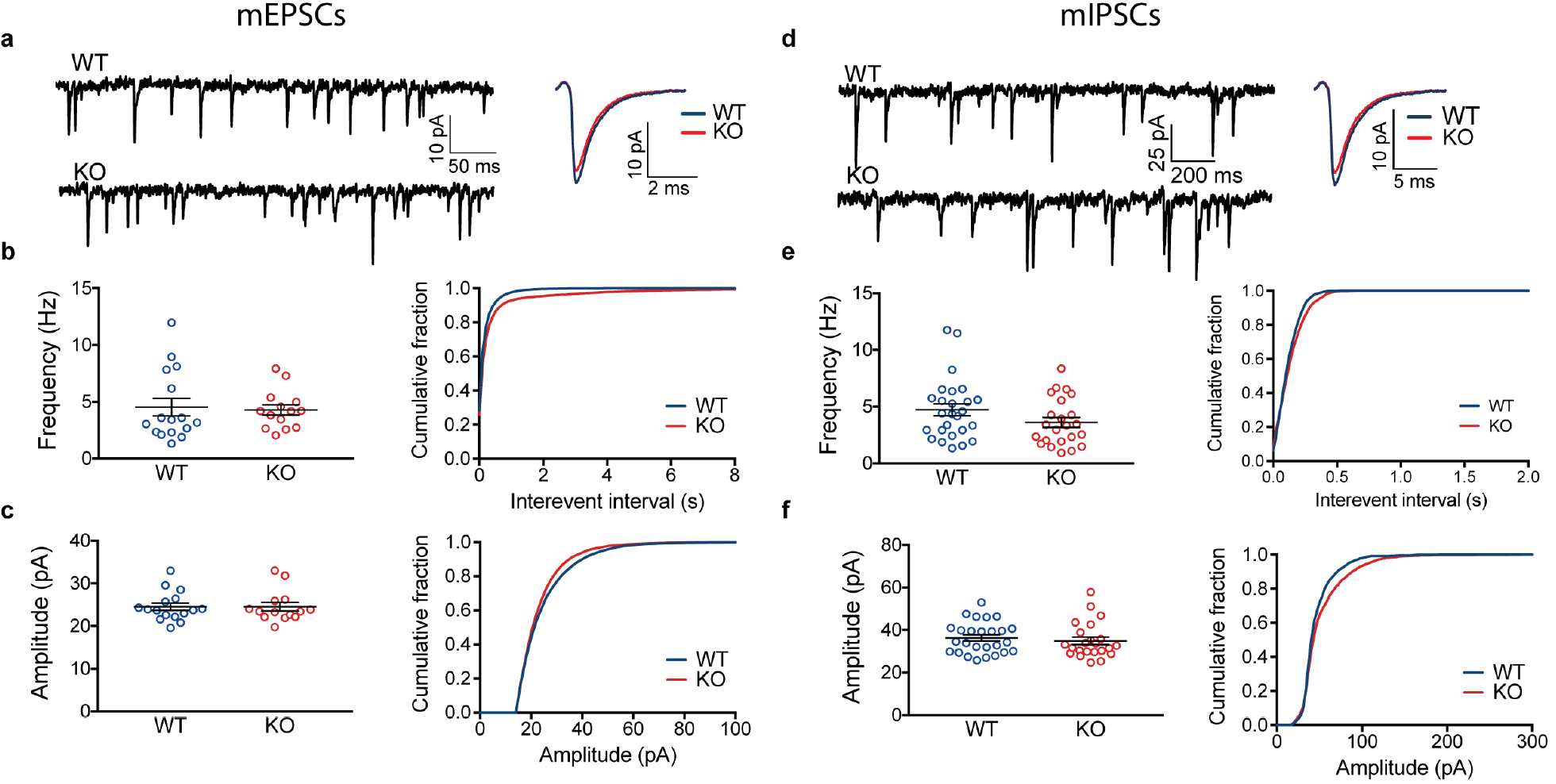
Cntnap2 KO mice show no significant alterations in PV+ inhibitory neuyron mEPSCs or mIPSCs. **a-c**, Frequency (WT 4.5 ± 0.8 Hz, KO 4.3 ± 0.5 Hz; *P* = 0.9901, Unpaired t test) and amplitude (WT, 24.5 ± 0.9 pA, KO 24.6 pA ± 1.0; *P* = 0.5970, Mann-Whitney test) of mEPSCs (WT *n* = 17, KO *n* = 15) and **d-f**, frequency (WT 4.7 ± 0.5 Hz, KO 3.6 ± 0.4 Hz; *P* = 0.4074, Mann-Whitney test) and amplitude (WT 36.0 ± 1.9 pA, KO 37.6 ± 2.5 pA; *P* = 0.8238, Mann-Whitney test) of mIPSCs (WT *n* = 28, KO *n* = 25) recorded from parvalbumin-positive (PV) inhibitory neurons are not statistically different between Cntnap2 KO and WT mice. Distribution of data is represented as box and whiskers plots with mean ± SEM.

**Supplementary Fig. 4.**
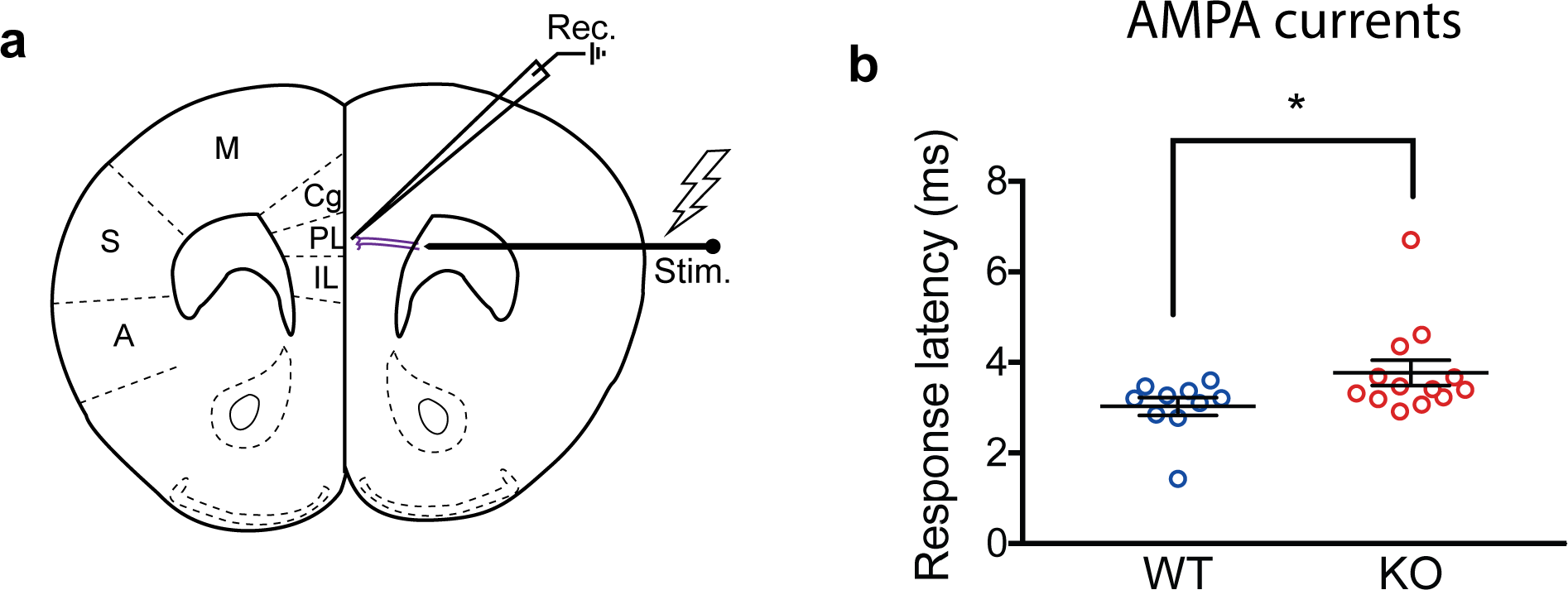
Increased stimulus response latency in L2/3 pyramidal neurons of Cntnap2 KO mice. **a**, Monopolar tungsten electrode was used to stimulate long-range axons (purple), which extend from the anterior forceps of the corpus callosum and project onto a patched excitatory neuron in L2/3 mPFC. **b,** Stimulus response onset for L2/3 pyramidal neurons in WT (3.03 ± 0.19 ms; *n* = 10 cells) and KO (3.77 ± 0.28 ms; *n* = 13 cells) mice shows an increase in evoked AMPA current response latency from time stimulus onset (\**P* = 0.0438, Mann-Whitney test).

**Supplementary Fig. 5.**
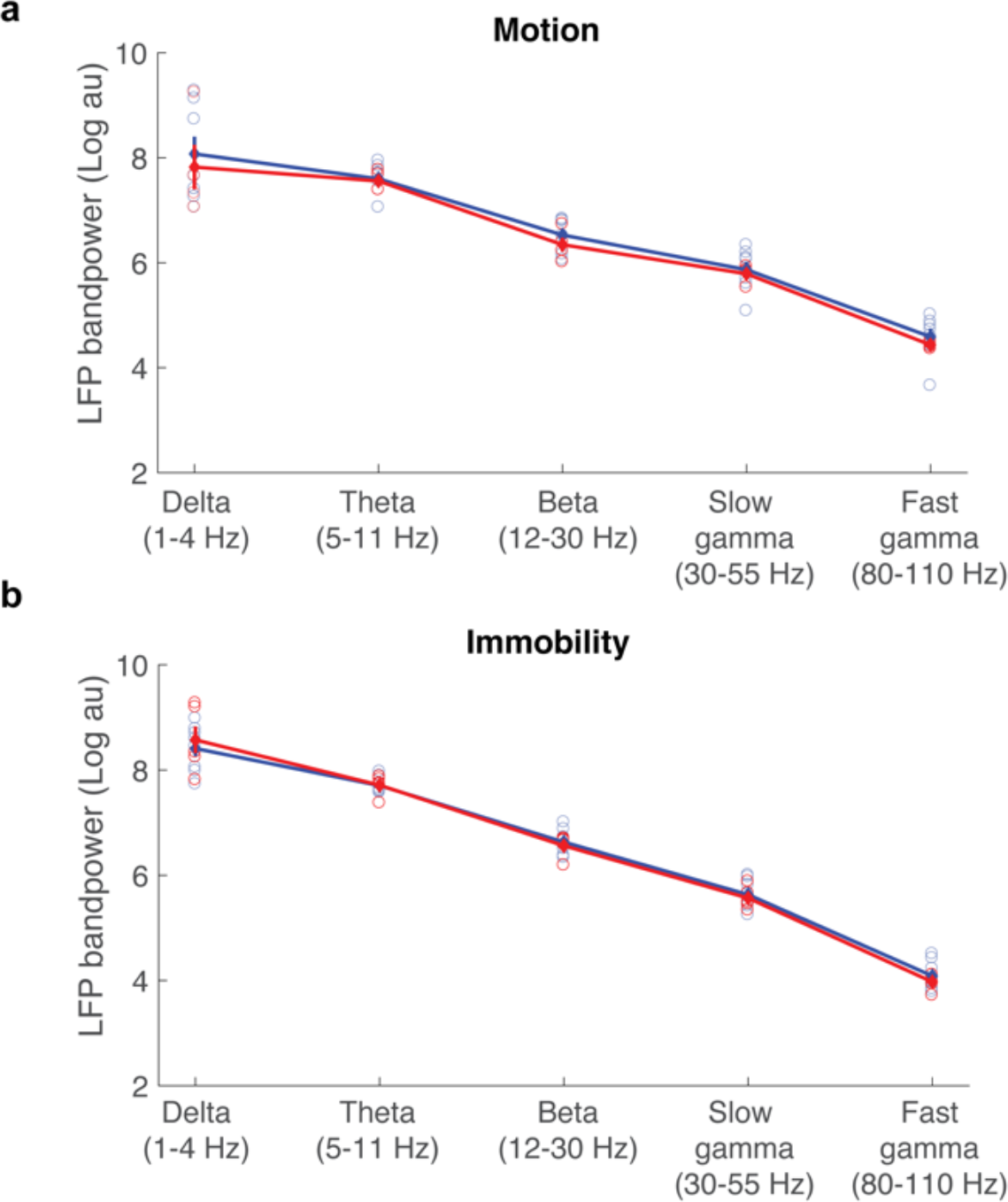
Local field potential power measurements in mPFC. Mean bandpower quantification of local field potential (LFP) during (**a**) motion and (**b**) immobility in WT and Cntnap2 KO mice. Power was calculated from a single representative channel per mouse, located in the prelimbic medial prefrontal cortex. No significant differences were observed (*P* > 0.05, Wilcoxon test).

**Supplementary Fig. 6.**
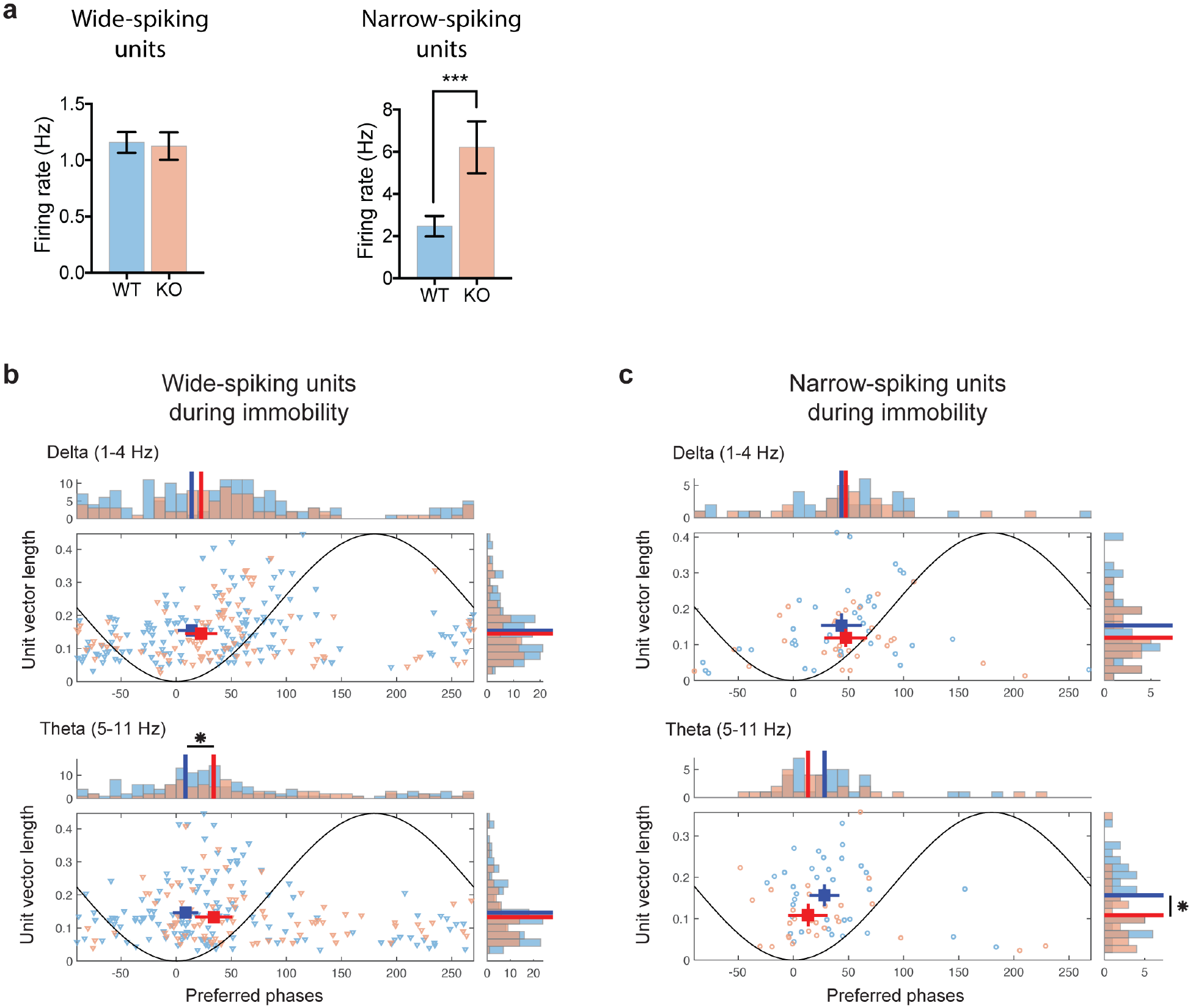
Firing rates, phase-locking, and phase-preference of identified wide-spiking (WS) and narrow-spiking (NS) units during immobility. **a**, Average firing rate during immobility per WS unit (top) and NS unit (bottom) in WT versus KO mice. WS units spiked with similar rates in both mouse groups, whereas NS units from KO mice fired significantly more than WT mice (*P* < 0.01; Wilcoxon test). **b-c**, Distributions and corresponding histograms of strength of phase-locking (mean vector length) versus preferred phases of pooled WS units (b) and NS units (c) from WT (blue) and KO mice (red). Plotted similarly to Figure 5f-g respectively. Preferred phases for WS units in KO animals are shifted to later phases, compared to WT. NS units from KO animals exhibit, on average, weaker phase locking than those of WT animals, in theta frequency LFP oscillations.

**Supplementary Fig. 7.**
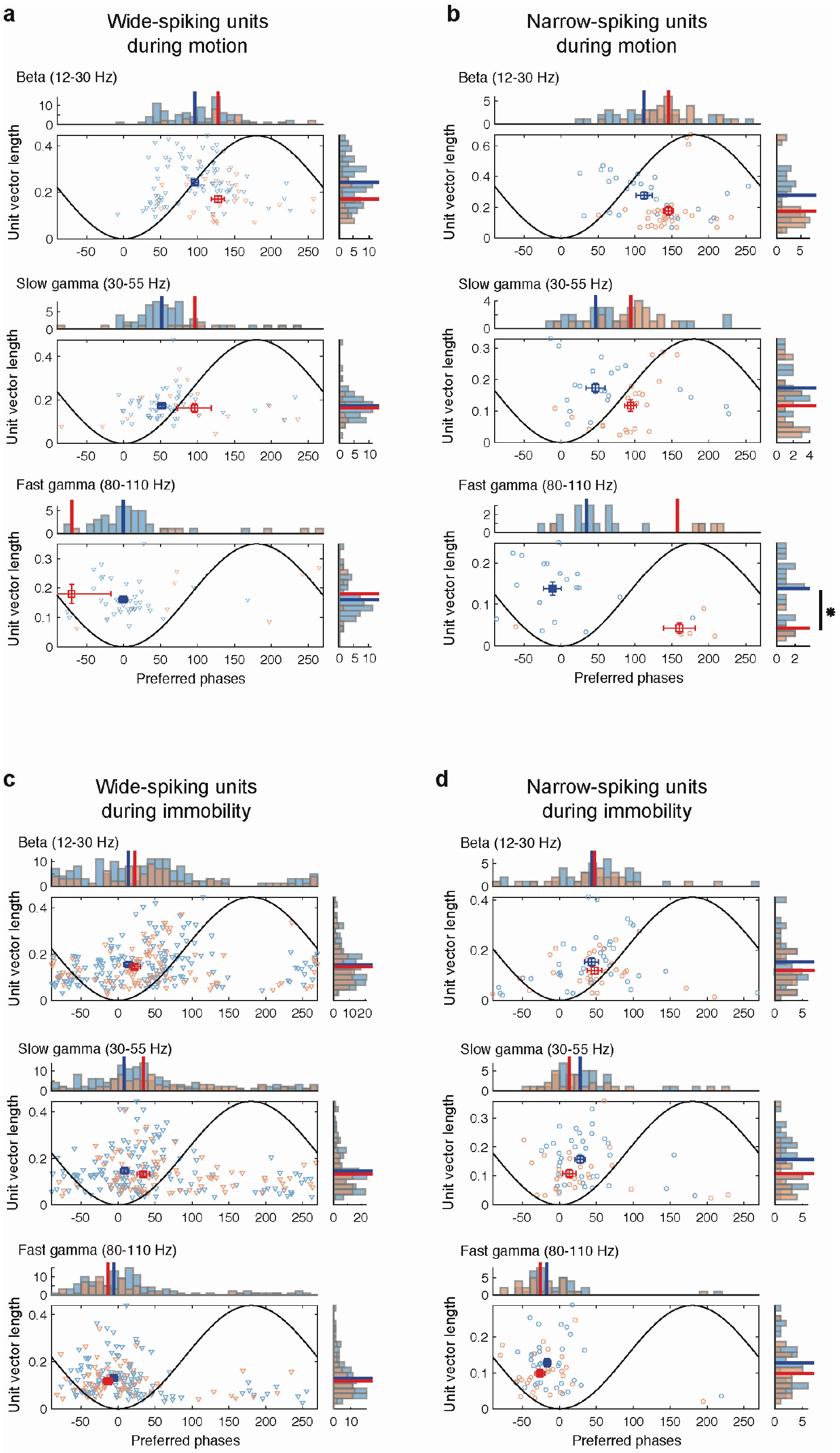
Phase-locking strength and phase-preference in identified WS and NS units for higher frequency oscillations. **a**,**c**, Distributions and corresponding histograms of strength of phase locking (mean vector length) versus preferred phases of pooled WS units from WT (blue) and KO mice (red) during motion (**a**) and immobility (**c**). Plotted similarly to Figure 5f-g and Supplemental Figure 5. Dashed lines indicate phase means of distributions that are not significantly non-uniform (*P* > 0.05; Rayleigh test for non-uniformity). **b**,**d**, Same as **a**,**c**, for pooled NS units of WT and KO mice.

**Supplementary Fig. 8.**
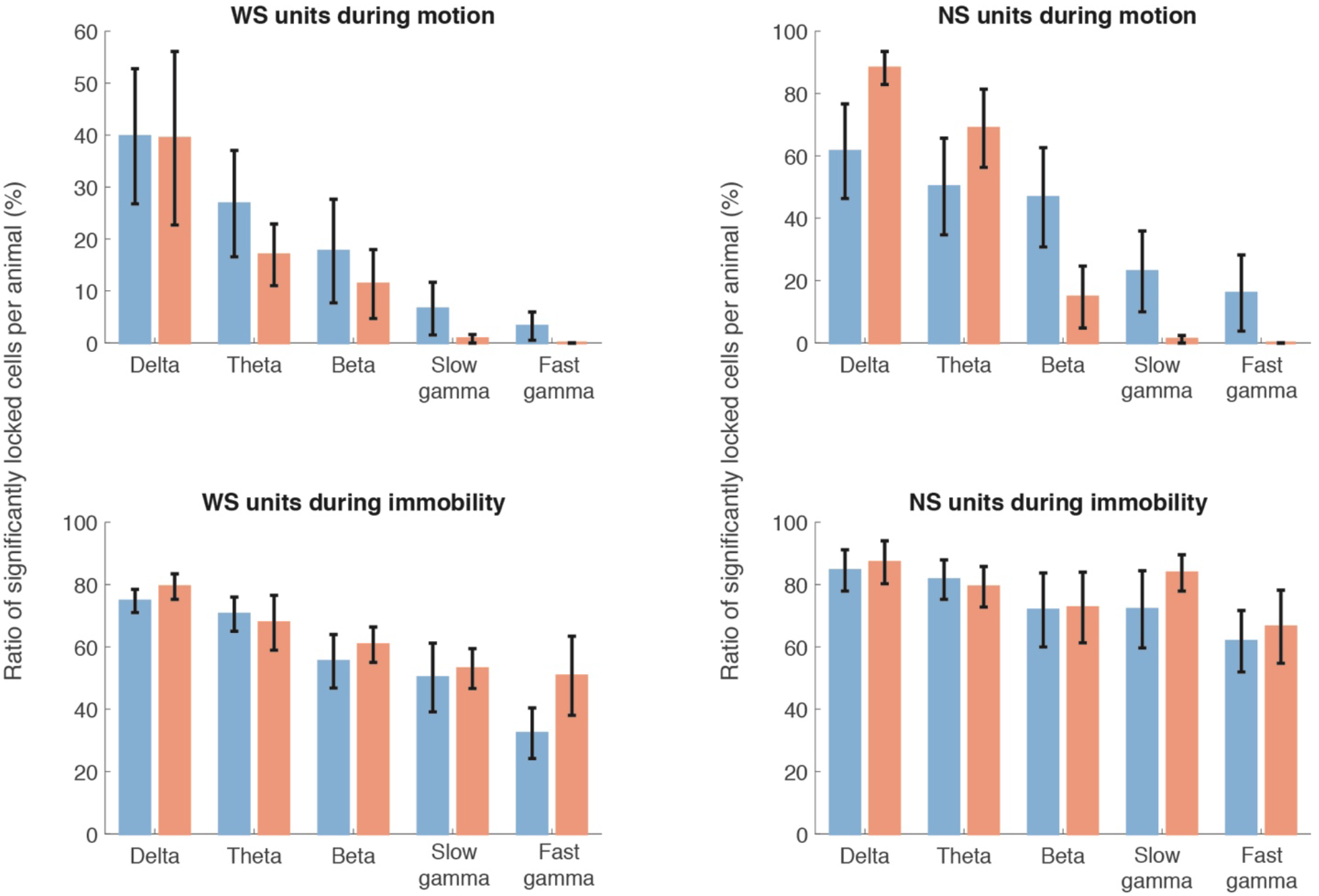
Ratio of phase locked cells. Mean percentage of WS units (left) or NS units (right) per animal, that were phase locked to each frequency range during motion (top) or immobility segments (bottom). Only cells that produced > 200 spikes were included in the analysis. No significant differences were observed between WT (blue) and KO animals (red). (*P* > 0.05, Wilcoxon test, Bonferroni corrected over all frequency range comparisons).

